# A bioengineered model of human placental exposure to environmental metals during pregnancy

**DOI:** 10.1101/2024.09.06.611636

**Authors:** Pouria Fattahi, Mousa Younesi, Won Dong Lee, Keumrai Whang, Taewook Kang, Joshua D. Rabinowitz, Lauren M. Aleksunes, Dan Dongeun Huh

## Abstract

Exposure of pregnant women to toxic metals is an environmental health issue associated with various pregnancy complications. Efforts to advance our biological understanding of this problem and mitigate its adverse effects, however, have been challenged by ethical concerns of human subject research during pregnancy. Here, we present an alternative approach that leverages the design flexibility, controllability, and scalability of bioengineered human reproductive tissues to enable experimental simulation and in-depth investigation of placental exposure to environmental metals in maternal circulation. Central to this method is an in vitro analog of the maternal-fetal interface and its dynamic tissue-specific environment constructed using primary human placental cells grown in a micro-engineered device. Using cadmium as a representative toxicant, we demonstrate the proof-of-concept of emulating the human placental barrier subjected to the flow of cadmium-containing maternal blood to show how this model can be used to examine adverse biological responses and impaired tissue function on both the maternal and fetal sides. Moreover, we present a mechanistic study of maternal-to-fetal cadmium transport in this system to reveal that efflux membrane transporters expressed by trophoblasts may play an important protective role against cadmium-induced toxicity. Finally, we describe metabolomic analysis of our microphysiological system to demonstrate the feasibility of discovering metabolic biomarkers that may potentially be useful for detection and monitoring of cadmium-induced placental dysfunction.

## Introduction

Pregnancy is an essential process of reproduction in which the female body undergoes remarkable remodeling in a highly coordinated fashion to support the development of the fetus^1–3^. By nature, most of the biological changes that occur during this process are transient, but altered maternal responses to external stimuli due to pregnancy can also result in lasting effects with long-term health consequences^4^. As a representative example, pregnancy is known to increase maternal sensitivity and susceptibility to adverse effects of foreign substances, especially those encountered in settings of environmental or occupational exposures^5^. Among such materials are toxic metals, such as lead, mercury, cadmium, and arsenic, which represent a class of environmental contaminants that can enter the maternal circulation through dermal exposure, ingestion, or breathing^6^. Epidemiology studies have shown that environmental metals are commonly found in pregnant women and that high-level exposures to these toxicants are associated with various pregnancy complications, including miscarriage, preterm birth, and low birth weight^7^. Many of these adverse outcomes have also been linked to impaired childhood development, as well as the increased risk of cardiovascular, neurological, and metabolic diseases in adult life^5,8,9^.

Despite widespread recognition of prenatal metal exposure as an important public health concern, much remains to be learned about the biological basis and consequences of this problem. Advances in environmental toxicology in the last decades have generated a wealth of information and insight on how metals can enter and accumulate in the human body and how metal-specific free radicals produced by the toxicants may elicit abnormal biological responses, including oxidative stress and cellular injury, tissue inflammation, disrupted protein homeostasis, and suppressed activities of essential enzymes^10^. Given that maternal physiology undergoes progressive, significant adaptations during gestation to accommodate fetal growth, however, it is questionable whether these general findings are directly translatable to understanding and predicting the negative health impact of environmental metal exposure in the specific context of pregnancy.

In particular, one of the key outstanding questions in this area is how environmental metals in the maternal system affect the placenta, which is arguably the least understood organ in the human body uniquely associated with pregnancy^11^. The placenta functions as the interface between the mother and the fetus that enables a controlled exchange of nutrients, oxygen, hormones, and other essential molecules, while also providing a restrictive barrier to prevent the passage of harmful substances from the maternal blood to fetal circulation^12^. Importantly, contrary to this general notion of the placenta as a protective barrier, researchers have revealed measurable levels of mercury, lead, arsenic, and cadmium in various clinical specimens of human pregnancy, including amniotic fluid^13,14^, cord blood^15,16^, and placenta explants^17^, showing the ability of some of these metals of maternal origin to cross the placental barrier and reach the fetal compartment. By demonstrating the potential of these materials as a threat to pregnancy, this body of work has provided a scientific basis for raising the poorly understood question of how toxic metals accumulating in fetal tissues affect the development of the fetus^6^.

Equally significant gaps exist, however, in our knowledge about the effect of toxic metals on the maternal-fetal interface of the placenta that plays a vital role in the maintenance of pregnancy. Specifically, it remains elusive whether the presence and trafficking of toxic metals can directly harm cellular components of the placental barrier. We also understand little about whether and how metal exposure can alter the biological phenotype and functional capacity of the placental barrier as an integrated tissue unit to dysregulate maternal-fetal communication and material exchange, which may have significant long-term implications in fetal development. Unfortunately, investigating these mechanistic questions in pregnant women continues to be a major challenge even though considerable progress has been made in clinical research of human pregnancy^18^. These types of studies are heavily restricted by ethical considerations and suffer from small cohort sizes of clinical studies and the difficulty of probing the delicate, inaccessible organ of pregnancy to measure and interrogate biological activities of its functional units in the native context.

As a step towards addressing this challenge, here we introduce a bioengineering approach for experimental modeling and in-depth preclinical studies of human placental exposure to environmental metals. This method is enabled by a microengineered system capable of using clinically sourced primary cells to generate vascularized human placental tissues that can be perfused in a controlled manner to recreate the maternal-fetal interface of the human placenta and its hemodynamic environment. We present a case study in which cadmium was used as an environmentally-relevant model toxicant to demonstrate how the bioengineered placental analog can be used to visualize and measure an array of biological responses and relevant functional endpoints to generate quantitative, high-content data useful for developing a more detailed understanding of metal toxicities. Through in-depth analysis of breast cancer resistance protein (BCRP) in this model, we also show the capacity of maternal efflux membrane transporters to reduce maternal-to-fetal transfer of cadmium and adverse biological responses of the placenta. Moreover, we demonstrate the potential of our platform for biomarker discovery by presenting unbiased global metabolomic analysis of the maternal and fetal compartments to identify tissue-specific metabolites that can indicate placental dysfunction due to cadmium toxicity.

## Results

### Design of bioengineered human placental tissue

In the placenta, maternal-fetal interactions occur through molecular exchanges between the maternal blood pooling in the intervillous space and the fetal blood circulating in the vasculature of the chorionic villi (**Fig. 1a**)^19^. Physically separating these two compartments is a multilayered tissue assembly known as the placental barrier that consists of trophoblast cells facing the maternal blood and the underlying connective tissue containing a network of capillary blood vessels and stromal cells (**Fig. 1b**). Our bioengineered placental tissue was designed to capture the multicellularity and 3D structural organization of this functional unit of the placenta that forms the maternal-fetal interface (**Fig. 1c**).

**Figure 1.**
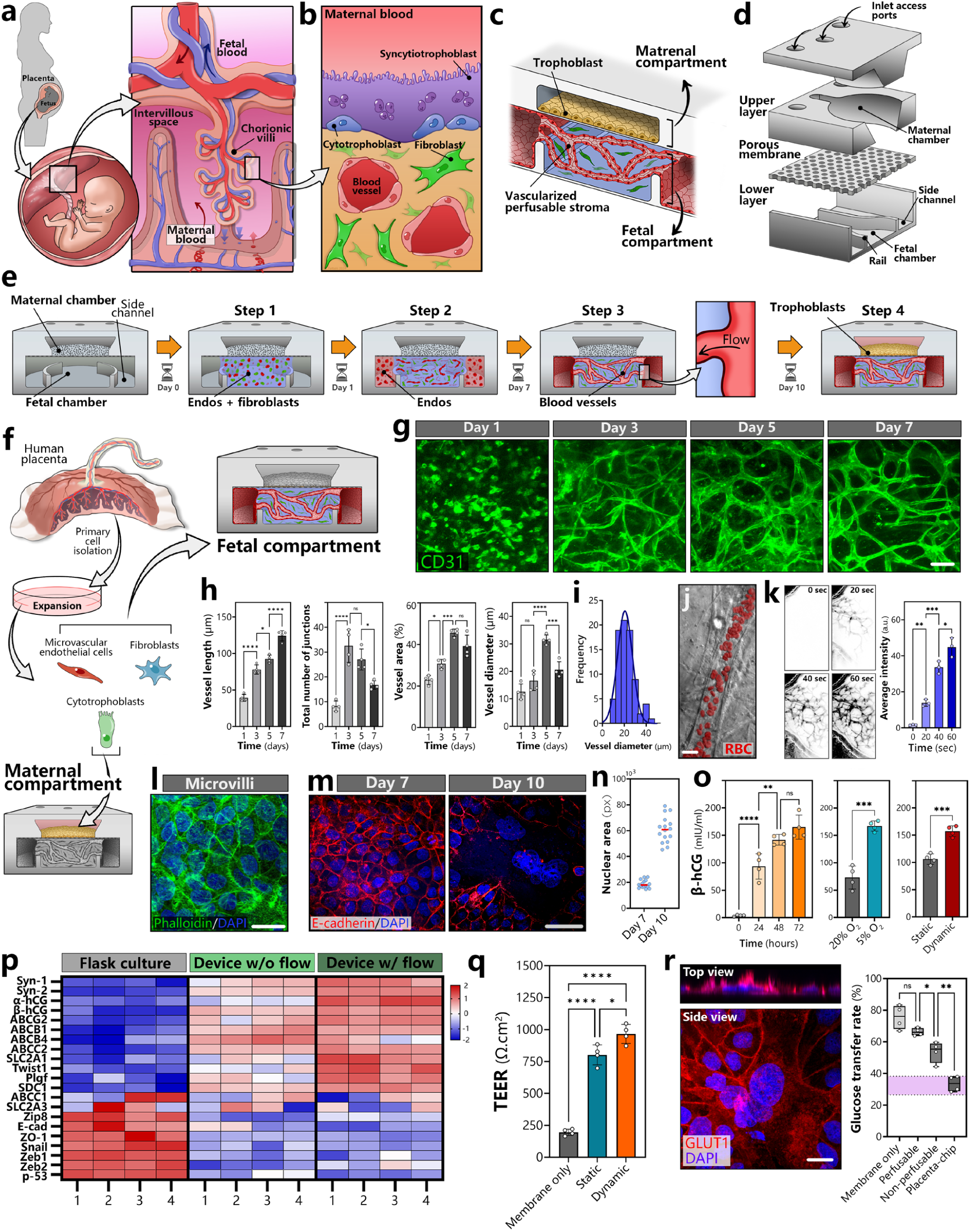
A bioengineered microphysiological model of the maternal-fetal interface in the human placenta. **a,b.** Schematics of the cotyledon in the human placenta (**a**) and the placental barrier (**b**). **c.** Illustration of a microengineered 3D model of the human placental barrier. **d.** Layer-by-layer view of the microdevice used to culture placental cells. **e.** A sequential process of human placental tissue production in the device. **f.** Illustration of primary human placental cells sourced from placenta explants for use in our model. **g.** Representative confocal images capturing the progression of vascular formation over a 7-day period. Scale bar, 100 μm. **h,i.** Quantification of select vascular features during vasculogenesis. **j,k.** Demonstration of vascular perfusion using red blood cells (RBCs) (**j**) and 70 kDa FITC-dextran (pseudo-colored black in the timelapse images) (**k**). Scale bar, 20μm. **l.** A confocal micrograph of microvilli shown with actin immunostaining (green). Scale bar, 25 μm. **m.** E-cadherin (red) and nuclear (blue) staining before (Day 7) and after (Day 10) spontaneous syncytialization. Scale bar, 50μm. **n.** Quantification of nuclear area in the trophoblast layer. **o.** Quantification of the level of β-human chorionic gonadotropin (β-hCG) during syncytialization over 72 hours from Day 7 through Day 10 (left), at different oxygen conditions (middle), and under static vs. dynamic culture conditions (right). **p.** A heatmap of gene expression analyzed by RT-PCR. **q.** Comparison of barrier integrity using transepithelial electrical resistance (TEER). **r.** Immunofluorescence of GLUT1 (left) and quantification of the rate of glucose transfer from the maternal to fetal chambers (right). Scale bar, 25μm. Data are presented as mean ± SD with *n* = 4. ns: not significant, *\*P < 0.05, **P < 0.01, ***P < 0.001,* and *\*\*\*\* P < 0.0001*.

This model is created in a multilayered device that consists of i) a 2-mm circular maternal chamber in the upper layer representing the mater blood-containing intervillous space, ii) another 2-mm circular fetal chamber in the lower layer that corresponds to the fetal compartment, and iii) a thin semipermeable membrane with 1-μm pores sandwiched between the two layers (**Fig. 1d**). The lower layer of the device also contains microchannels on either side of the fetal chamber, and these two compartments are separated by a micropatterned rail structure protruding from the floor of the device (**Fig. 1d**). Each chamber is connected to its own set of access ports contained in another substrate bonded to the upper layer to permit independent control of cells and their fluidic environment, as well as sampling of device effluent in a compartment-specific manner (**Fig. 1d**).

The first step of engineering placental tissues in this device is to inject an extracellular matrix (ECM) hydrogel precursor mixed with human placental endothelial cells and fibroblasts into the fetal chamber and induce gelation to form a cell-laden hydrogel construct (**Step 1** in **Fig. 1e**). Next, another batch of placental endothelial cells is seeded into the side channels adjacent to the fetal chamber and grown on the channel surfaces (**Step 2** in **Fig. 1e**). In this configuration, the endothelial cells cultured in the hydrogel undergo a dynamic process reminiscent of vasculogenesis in vivo in which they become elongated and connect with one another to self-assemble into a 3D network of open blood vessels (**Step 3** in **Fig. 1e**). During this process, the endothelial lining of the side channels sprouts into the hydrogel and anastomoses with the 3D vessels, making the self-assembled vascular network accessible and perfusable from the side channels. Finally, human trophoblast cells are seeded into the maternal chamber and grown on the upper side of the intervening membrane to form a confluent monolayer (**Step 4** in **Fig. 1e**).

### Production and characterization of bioengineered human placental tissue

To model the human maternal-fetal interface with high biological fidelity, we used primary cytotrophoblasts, villous microvascular endothelial cells, and fibroblasts isolated from term human placentas after delivery (**Fig. 1f**). Importantly, given the sensitivity of primary human placental cells to oxygen^20^, these cells were maintained at 5% oxygen, which approximates physiological oxygen levels in the human placenta^21,22^, throughout culture to allow them to preserve their native cellular phenotype.

For tissue production in the device, we first prepared a mixture solution containing placental endothelial cells, fibroblasts, fibrinogen, and thrombin, and injected it into the fetal chamber. During this step, the solution was stably pinned along the microfabricated rails due to surface tension, and the advancing meniscus was guided by this structure until the entire chamber was filled (**Supplementary** Fig. 1). This design permitted spatial confinement of the injected mixture in the fetal chamber without leakage to the side channels and the maternal chamber. Fluorescence microscopy of the endothelial cells during 3D culture in the fibrin scaffold revealed rapid reorganization of individual cells into interconnected endothelial tubes, which was visible within 3-4 days of culture (**Fig. 1g**). The self-assembled blood vessels appeared to stabilize over time, yielding an average diameter of 19.9 μm by day 7 (**Figs. 1h, 1i**).

The vasculature also became perfusable within 7 days in this model (**Figs. 1j, 1k**), after which the entire network was constantly perfused with media throughout the culture period. Interestingly, the blood vessels maintained in this flow condition appeared morphologically different from those in static conditions without vascular perfusion. The most notable difference was the smaller size of these vessels (**Supplementary Fig. 2a**), which might be viewed as a better representation of their microvascular origin. This observation was verified by quantitative comparison of vascular features that showed statistically significant reduction in vessel diameter, area, and the number of junctions in the dynamic culture condition with flow (**Supplementary Fig. 2a**).

In the upper half of the device, cytotrophoblasts were introduced into the maternal chamber on day 7 after the completion of vasculogenesis in the underlying fetal chamber. With the entire system maintained at 5% O2 while being perfused with media, these cells remained proliferative and continued to grow for 3-4 days to form a confluent monolayer covered with microvilli on the apical surface (**Fig. 1l**). Interestingly, the density of the microvilli was significantly lower when the trophoblasts were cultured in the absence of flow (**Supplementary Fig. 2b**). Over the next 3 days, the trophoblast cells in the monolayer spontaneously fused and began to exhibit the morphological characteristics of the placental syncytium in vivo, which was evidenced by a loss of intercellular junctions and aggregation of cell nuclei (**Figs. 1m, 1n**). This physiological differentiation of cytotrophoblasts into syncytiotrophoblasts was observed in the monolayer surface and further verified by a progressive increase in the level of β–human chorionic gonadotropin (β-hCG) in the effluent of the maternal chamber over time (**Fig. 1o**). Of note, our data revealed significantly higher levels of β-hCG due to the lower oxygen tension (5%) and flow conditions used in our model (**Fig. 1o**), illustrating the capacity of the physiologically relevant environment to help the cultured trophoblasts express more differentiated phenotype.

For further characterization of the syncytium, we then conducted RT-PCR to measure and compare the expression of placental cell markers. Analysis of the syncytialized cells isolated from our model showed marked upregulation of a set of genes with known associations with syncytiotrophoblast differentiation, such as syncytin, β-hCG, SLC2A1, and PLGF, when compared to trophoblasts cultured in flasks or our device under static conditions without flow for the same amount of time (**Fig. 1p**). In comparison to flask culture, our engineered placental model was also seen with decreased expression of intercellular junctional proteins (e.g., E-cad, ZO-1) and epithelial-mesenchymal transition markers such as Zeb2, whose expression levels are known to decrease with placental maturation^23^. Similar differences were observed in the comparison of the same markers between flask culture and the engineered model without flow but in this case, the extent of changes, especially increases in the expression of syncytiotrophoblast differentiation markers, was not as substantial (**Fig. 1p**).

This enhanced differentiation of the maternal-fetal interface in our model resulted in improved functional capacity as a restrictive barrier, which was demonstrated by increased levels of electrical resistance between the maternal and fetal chambers (**Fig. 1q**). Expression of glucose transporter 1 (GLUT1) – the most abundant type of glucose transporters in the human placental barrier^24^ – was another functional marker measured in our analysis, which was shown by robust immunofluorescence staining along the apical surface of the trophoblast layer (**Fig. 1r**). Importantly, temporal measurement of glucose concentrations in the maternal and fetal perfusate yielded an average maternal-to-fetal transfer rate of 33.4 %, which is within the physiological range of glucose transfer rates measured in perfused human placenta explants^25,26^ (**Fig. 1r**).

### Modeling and analysis of cadmium toxicity in the human placenta

Next, we explored the utility of our bioengineered model for in-depth in vitro investigation of how environmental toxicants affect the maternal-fetal interface of the human placenta. This study focused on modeling placental toxicity of cadmium, an environmental metal monitored as a high-priority toxicant by environmental agencies due to its known or suspected adverse effects on human health^27,28^. Cadmium is commonly found in the environment, and humans are exposed to this substance in a variety of situations, including food consumption, drinking water contamination, metal mining, fertilizer production, fossil fuel combustion, and tobacco smoking^29^. Human exposure to cadmium occurs globally, but its prevalence has been particularly high in developing countries, which makes it an important global health equity issue^30^.

Notably, cadmium is known to pose significant threats to pregnancy. According to a recent systematic review of epidemiological literature, for example, cadmium exposure increases the risk of low birth weight and preterm birth by 21% and 32%, respectively.^31^ Studies have also reported measurable levels of cadmium in human placentas and umbilical cord blood collected after delivery^32^. Based on this evidence, it is suspected that cadmium accumulating in the placenta may exert direct or indirect deleterious effects on the maternal-fetal interface and its function, playing a causative role in the development of adverse pregnancy outcomes.

The biological mechanism of how this process of placental dysregulation can occur, however, remains poorly understood. Although reproductive toxicity of cadmium has been investigated in various animal species^33,34^, human data are extremely limited, hampering our fundamental understanding of this problem. Therefore, our goal was to use our bioengineered tissue as a platform to model human placental exposure to cadmium, modulate its key parameters in a precisely controlled manner, and generate informative human-relevant data.

To this end, we established an exposure model (**Fig. 2a**) by adding cadmium chloride to the flow of trophoblast media in the maternal chamber. This model was first used to assess acute injury of the placental barrier after 24 hours of treatment in a range of cadmium concentrations previously described in the literature. At lower doses ranging from 0.1 to 10 μM, which are more relevant to human situations^35^, direct visualization of the trophoblast layer using live/dead staining showed intact or minimally affected cells, but treatment with higher concentrations (50-100 μM) resulted in larger numbers of membrane-compromised cells (**Fig. 2b**). Interestingly, at a given dose, the extent of this cellular injury was substantially greater in a Transwell-based exposure model constructed by growing the same batches of primary human trophoblasts and villous endothelial cells on either side of the Transwell membrane for the same duration (**Fig. 2c**), which was used to represent the gold standard model for in vitro studies of cadmium toxicity^36,37^.

**Figure 2.**
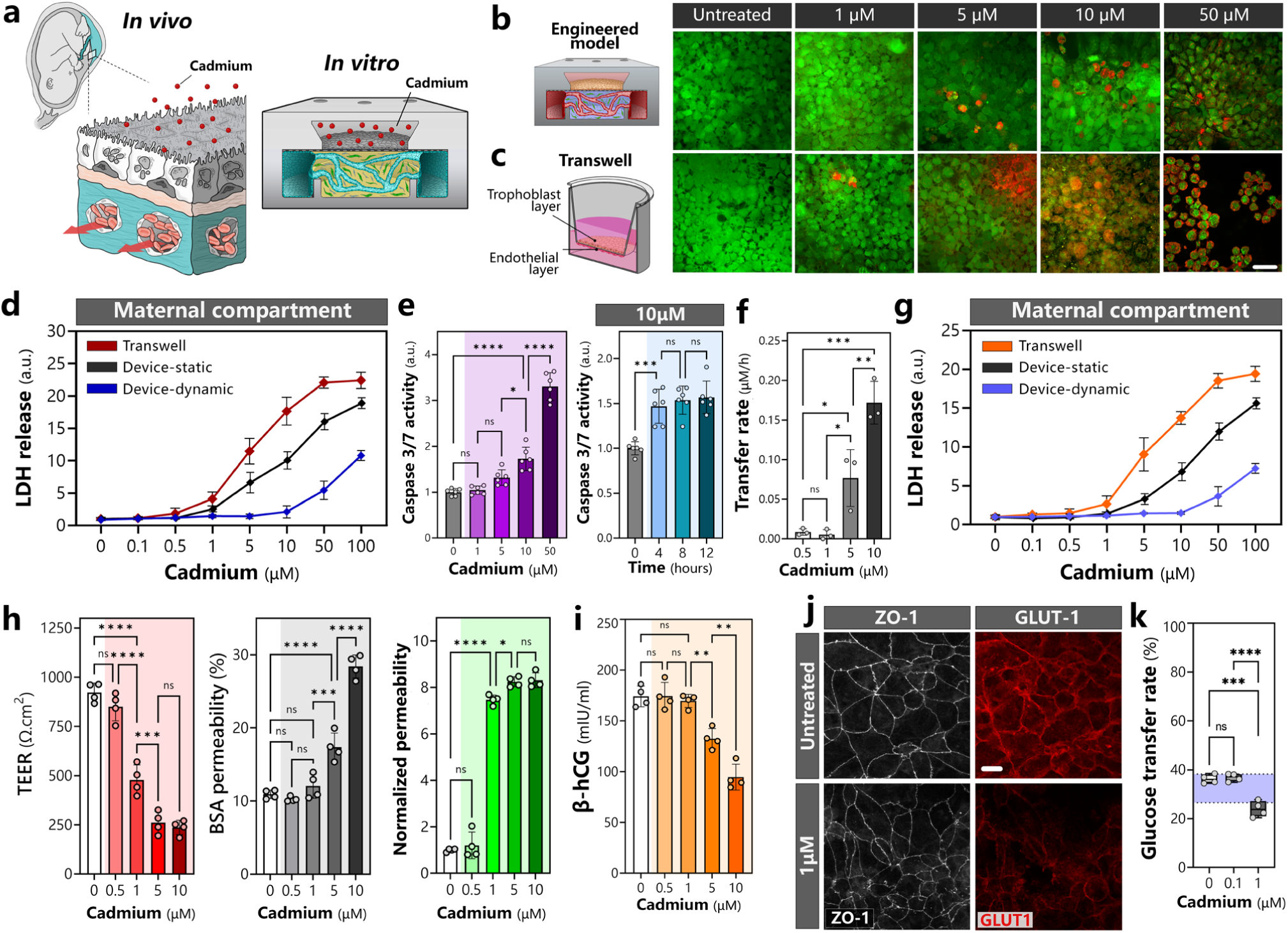
Simulation of cadmium exposure in the engineered placental model. **a.** Schematic illustrations of in vitro techniques to model exposure of the in vivo human placental barrier to cadmium using our platform. **b,c.** Fluorescence micrographs of trophoblasts exposed to varying concentrations of cadmium in the engineered model (**b**) and Transwell (**c**). Green and red fluorescence show live and membrane-compromised cells, respectively. Scale bar, 20 μm. **d.** Quantification of LDH release from the maternal compartment after 24 hours of cadmium exposure. ‘Device-static’ and ‘Device-dynamic’ represent cadmium exposure in the engineered model in the presence and absence of flow, respectively. Data are normalized to the untreated control group (0 µM). **e.** Plots of caspase-3/7 activities after 24 hours of exposure to varying cadmium concentrations (left) and at 10 µM (right). **f.** Quantification of maternal-to-fetal cadmium transfer. **g.** Quantification of LDH release from the fetal compartment after 24 hours of cadmium exposure. ‘Device-static’ and ‘Device-dynamic’ represent cadmium exposure in the engineered model in the presence and absence of flow, respectively. Data are normalized to the untreated control group (0 µM). **h.** Effects of cadmium on barrier function as assessed by TEER (left), permeability to fluorescently labeled bovine serum albumin (BSA) (middle), and permeability to lucifer yellow (LY) dye (right). LY data were normalized to the untreated control group (0 µM). **i.** Quantification of β-hCG secretion by trophoblasts in the maternal chamber. **j.** Immunofluorescence imaging of ZO-1 (left column) and GLUT1 (right column) transporters expressed by trophoblasts. Scale bar, 20 μm. **k.** Evaluation of the rate of maternal-to-fetal glucose transfer. Data are presented as mean ± SD with *n* = 4. ns = not significant, *\*P < 0.05*, *\*\*P < 0.01*, *\*\*\*P < 0.001*, and *\*\*\*\* P < 0.0001*.

Consistent with these observations, the concentrations of lactate dehydrogenase (LDH) in the effluent of the maternal compartment exposed to lower doses of cadmium were statistically indistinguishable from those measured in the untreated group, and appreciable increases from the baseline levels were only detected at 10 μM or higher (**Fig. 2d**). By comparison, in the Transwell model and the engineered tissue maintained under static conditions without flow, the maternal LDH levels began to increase substantially even at 1 μM and were found to be more than twice as high as those in the dynamic model with flow at any given cadmium concentration greater than 1 μM (**Fig. 2d**). Our data also revealed significant upregulation of caspase activities in the trophoblast layer within a few hours of treatment at 10 and 50 μM of cadmium (**Fig. 2e**), indicating the potential of high-level cadmium exposure to induce rapid activation of apoptosis signaling in the placental syncytium.

The observation of this concentration-dependent cytotoxicity in the maternal compartment led us to examine the effects of maternal cadmium on the fetal side of our model. For this study, we first analyzed maternal-to-fetal transfer of cadmium by collecting device effluent from both the maternal and fetal chambers and measuring cadmium concentrations in the collected samples using nanoengineered plasmonic sensors designed specifically for cadmium detection and quantification (**Supplementary Fig. 3**). In all conditions tested, this analysis revealed the presence of cadmium in the fetal compartment within 24 hours of exposure, but the rate of transfer was dependent upon the cadmium level on the maternal side. At a cadmium chloride concentration of 1 μM, the transfer rate was very low (0.004 μM/h), consistent with the limited maternal-to-fetal transfer in human pregnancies (**Fig. 2f**). At 10 μM, however, approximately 47.3% of maternal cadmium was transferred to the fetal compartment at a rate of 0.17 μM/h. Notably, cadmium in the fetal compartment was found to induce LDH release from the vascularized stroma in concentration-dependent manners similar to those observed on the maternal side, but the threshold level of cadmium at which tissue injury became evident in our dynamic model was higher (50 μM) (**Fig. 2g**).

As expected, these cytotoxic effects produced by higher levels of cadmium also had a negative impact on the functional capacity of the engineered placental barrier. For example, at 10 μM, our exposure model showed compromised barrier function as demonstrated by significantly reduced electrical resistance across the barrier and its increased permeability to indicator dye molecules (**Fig. 2h**). What was unexpected, however, was substantially increased barrier permeability even at 5 μM (**Fig. 2h**), which was not associated with elevated LDH secretion and caspase signaling (**Figs. 2d, 2e, 2g**). Cadmium exposure at this concentration also resulted in significant decreases in the production of β-hCG (**Fig. 2i**), suggesting that lower, non-cytotoxic levels of cadmium may still generate adverse effects on the placental barrier.

We then asked whether and how cadmium exposure affects the essential physiological function of the placental barrier as a mediator of maternal-to-fetal glucose transfer. Since our results showed compromised, leaky barrier at 5 μM and higher, we tested lower cadmium concentrations between 0 and 1 μM. When treated with 0.1 or 1 μM for 24 hours, the syncytialized trophoblast layer remained structurally intact and did not show any signs of a loss of barrier integrity (e.g., intercellular gaps). At 1 μM, however, the cells were seen with noticeable changes in the expression of glucose transporters, which was best characterized by decreased surface area of the barrier stained positive for GLUT1 (**Fig. 2j**). Supporting these results, the measurement of glucose levels in the maternal and fetal compartments revealed significantly decreased glucose transfer rates, which fell below the physiological range (**Fig. 2k**). This adverse response was not observed when the model was exposed to 0.1 μM.

### Cadmium-induced inflammation and disruption of placental signaling

Clinical studies of prematurity have shown that inflammation in the placenta is associated with several clinical markers of fetal growth restriction and may play an important role in preterm birth^38^. Based on this clinical evidence and the known association of prenatal cadmium exposure with low birth weight and preterm birth^39^, we reasoned that cadmium in the maternal blood may induce inflammation of the placental barrier and that this adverse response may be one of the ways in which cadmium exposure can disrupt placental signaling that may eventually lead to adverse pregnancy outcomes reported in the literature.

To investigate relevant biological responses in our exposure model, we measured the levels of four representative pro-inflammatory cytokines, including interleukin-8 (IL-8), IL-6, IL1β, and tumor necrosis factor-alpha (TNF-α), all of which have been implicated in human placental inflammation^40^. Analysis of maternal effluent following 24 hours of exposure showed dose-dependent inflammatory responses in which all four cytokines were produced in significantly increased amounts when the model was subjected to 5 μM or higher (**Fig. 3a**), which coincided with the threshold above which barrier function and β-hCG production were compromised (**Figs. 2h, 2i**). Treatment with lower doses (0.5, 1 μM) yielded no differences in the cytokine levels (**Fig. 3a**). Similar to our observation of platform-dependent susceptibility to cadmium-induced tissue damage (**Figs. 2b-d**), the levels of the same cytokines were 1.5 – 3-fold higher in the Transwell model (**Fig. 3a**).

**Figure 3.**
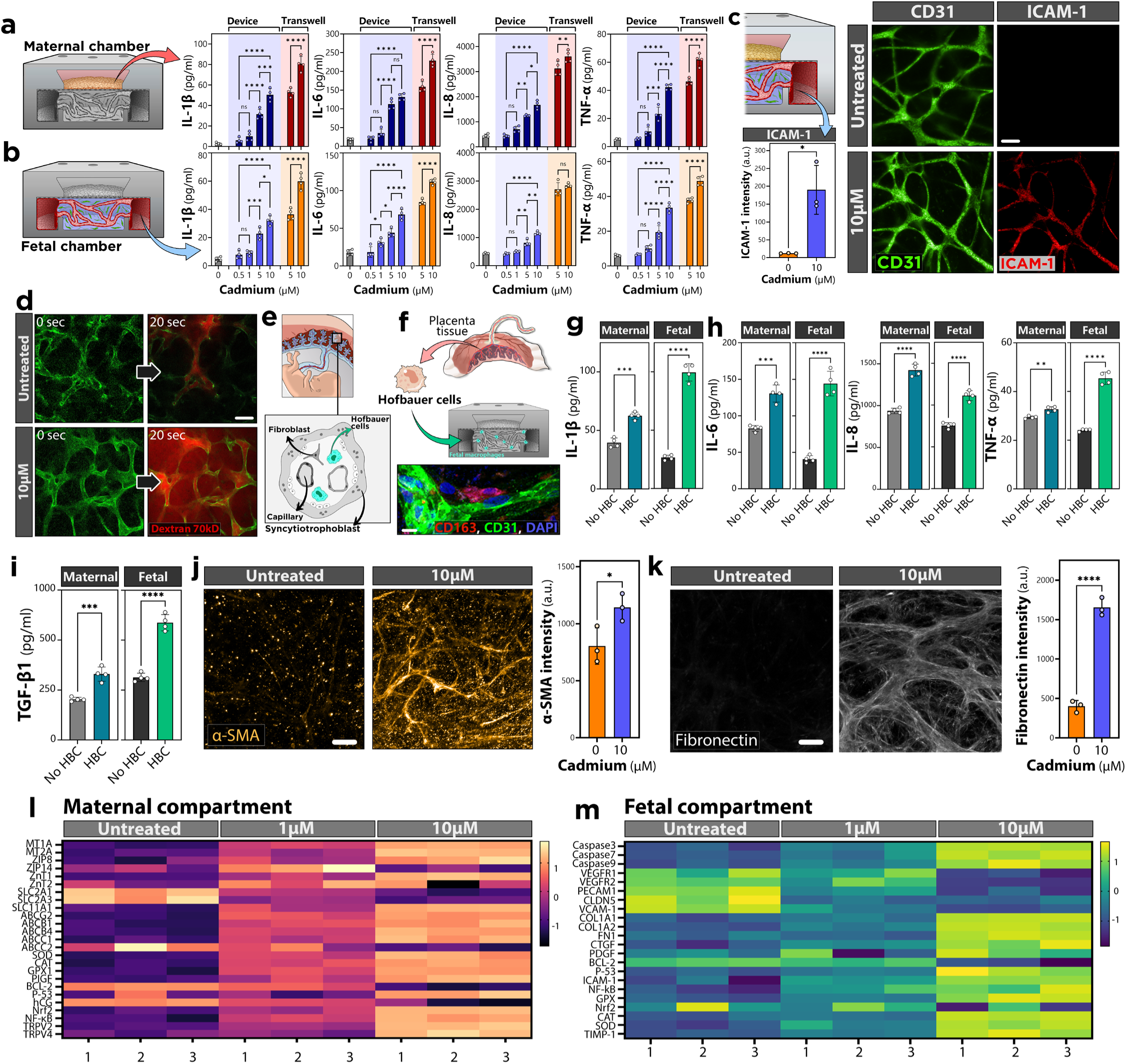
Responses of engineered placental barrier to cadmium. **a,b.** Analysis of select pro-inflammatory cytokines in media effluent collected from the maternal (**a**) and fetal (**b**) compartments of our model (labeled “Device”) and Transwell. **c,d.** Comparison of ICAM-1 expression in the vasculature (**c**) and vascular permeability (**d**). Scale bars, 100 μm. **e.** Schematic illustration of Hofbauer cells present in the maternal-fetal interface in vivo. **f.** Immunofluorescence imaging of CD163+ primary human Hofbauer cells incorporated into the engineered placental barrier model and grown for seven days. Scale bar, 20 μm. **g-i,** Evaluation and comparison of cytokine production in the presence or absence of Hofbauer cells (HBC). **j,k.** Increased expression of α-SMA (**j**) and fibronectin (**k**) due to cadmium exposure. Scale bars, 100 μm. **l,m.** Heatmaps illustrating altered gene expression in the maternal (**l**) and fetal (**m**) compartments due to cadmium analyzed by RT-PCR. Data are presented as mean ± SD with *n* = 4. Ns = not significant, *\*P < 0.05, **P < 0.01, ***P < 0.001,* and *\*\*\*\* P < 0.0001*.

For each cytokine, measurements taken from the effluent of the fetal compartment of our model showed the same concentration-dependent trends, as well as lower cytokine levels than the Transwell control (**Fig. 3b**). The pro-inflammatory microenvironment of the fetal chamber resulting from higher-level cadmium exposure also led to the activation of the vasculature in the stroma as illustrated by increased endothelial expression of intercellular adhesion molecule-1 (ICAM-1) at 10 μM (**Fig. 3c**). The inflamed blood vessels appeared to retain their architecture but were found to be leakier during perfusion with 70 kDa FITC-dextran (**Fig. 3d**), indicating cadmium-induced vascular permeability.

To more faithfully model the inflammatory milieu of the cadmium-exposed placental barrier, we then obtained donor-matched primary human Hofbauer cells, which are placental macrophages of fetal origin that reside in the villous stroma^41,42^ (**Fig. 3e**), and incorporated them into the fetal compartment of our system (**Fig. 3f**). During tissue production, these cells were found mainly in the vicinity of the self-assembled blood vessels (**Fig. 3f**). Notably, when the Hofbauer cell-containing model was treated with maternal cadmium at 10 μM for 24 hours, the production of IL-1β in the effluent of the maternal and fetal chambers was increased by 1.58 and 3.75 folds, respectively, as compared to control without Hofbauer cells (**Fig. 3g**). Similarly significant increases were observed in the analysis of the other cytokines (**Fig. 3h**), demonstrating the pro-inflammatory phenotype of Hofbauer cells previously suggested by histological studies of human placentas with inflammatory conditions (e.g., villitis)^43–46^.

Interestingly, inclusion of Hofbauer cells in the stromal compartment also resulted in substantially elevated levels of transforming growth factor-beta (TGF-β1) produced by cadmium exposure (**Fig. 3i**). Given previous studies showing the association of activated TGF-β1 signaling with fibrosis in preeclamptic human placentas^47^, this result prompted us to examine the fibroblast population in our exposure model. Immunofluorescence analysis of alpha-smooth muscle actin (α-SMA) revealed that the level of placental fibroblast activation was significantly increased as a results of treatment with 10 μM of cadmium chloride (**Fig. 3j**). Further supporting this observation, the stroma of the exposed model was also seen with profibrotic matrix remodeling as shown by increased deposition of fibronectin selected as a representative ECM (**Fig. 3k**), suggesting the capacity of cadmium to induce fibrogenic responses in the placental barrier.

Finally, we performed RT-PCR analysis of our model with the goal of gaining further insight into how placental signaling may be altered or dysregulated by cadmium exposure. Examination of top 25 differentially regulated genes in the maternal compartment showed increased activities of biological signaling involved in cellular responses to oxidative stress, which was evident from significantly elevated expression of metallothionein (MT), glutathione perioxidase-1 (GPX1), superoxide dismutase (SOD), catalase (CAT), and transcription factor nuclear factor erythroid 2-related factor 2 (Nrf2) (**Fig. 3l**). NF-κB was another gene related to oxidative stress and inflammation that was upregulated in the cadmium-treated model (**Fig. 3l**). Data also revealed signaling changes indicative of disrupted placental function, including dose-dependent reduction in the expression of hCG, as well as solute-carrier (SLC) genes, such as SLC2A1 and SLC2A3, that are responsible for encoding the glucose transporters^48^ (**Fig. 3l**). This result was in agreement with the findings of β-hCG ELISA (**Fig. 2i**) and immunofluorescence of GLUT1 (**Fig. 2j**). On a related note, cadmium exposure upregulated a set of genes that encode ATP-binding cassette (ABC) transporters known to play an important role in regulating transport function of the placental barrier^49^ (**Fig. 3l**).

PCR analysis of cells collected from the fetal compartment of the cadmium-treated model showed increased signaling that mediates apoptosis (e.g., caspase-3, caspase-7) and antioxidant responses (e.g., SOD, CAT, GPX, Nrf2, NF-κB) (**Fig. 3m**). Consistent with our observation of cadmium-induced vascular inflammation (**Fig. 3c**) and fibrogenesis in the villous stroma (**Figs. 3j, 3k**), upregulated genes in the exposure model also included ICAM-1, COL1A1, COL1A2, FN1, and CTGF (**Fig. 3m**). Data indicated that cadmium treatment resulted in dose-dependent decreases in the expression of vascular endothelial growth factor receptor (VEGFR) genes involved in placental angiogenesis^50^ (**Fig. 3m**). Also included in this group of downregulated genes was claudin-5 (CLDN5) that plays an important role in maintaining placental barrier function, especially in regulating ion/electrolyte transport^51^.

### Investigation of placental efflux transporters

One of the findings of our PCR analysis that was particularly interesting was the increased expression of ABC genes in the trophoblast cells due to cadmium exposure (**Fig. 3l**). This subset included ABCG2, ABCB1, ABCB4, and ABCC1 that are known to encode breast cancer resistance protein (BCRP), multidrug resistance protein 1 (MDR1), MDR3, and multidrug resistance-associated protein 1 (MRP1), respectively. Notably, these are transmembrane proteins expressed by syncytiotrophoblasts in the human placenta that utilize ATP-derived energy to actively transport substances from the intracellular compartment to the extracellular space to prevent fetal exposure to xenobiotics^52,53^ (**Fig. 4a**). Although the general function of these proteins as efflux transporters has been established, their specific role and significance in placental toxicity of cadmium remain to be further investigated.

**Figure 4.**
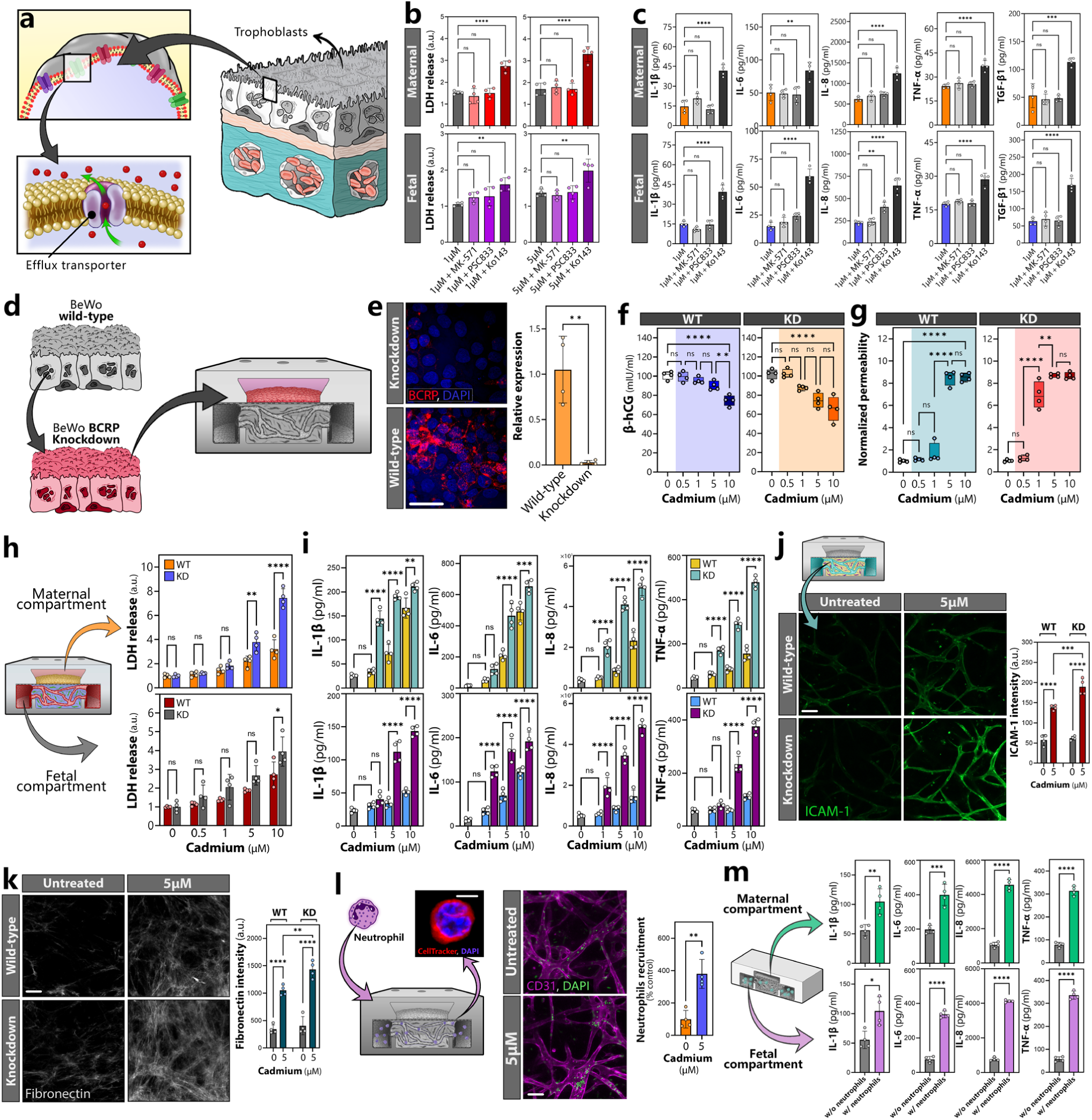
Investigation of BCRP in the cadmium exposure model. **a.** Schematics of efflux transporters expressed on the maternal blood-facing apical surface of trophoblasts in the human placental barrier. **b,c.** Quantification of cadmium-induced LDH release (**b**) and pro-inflammatory cytokine production (**c**) when efflux transporters are biochemically inhibited. Data in (**b**) were normalized to the untreated control group (not shown in the plot). **d.** Illustration of incorporating BeWo cells with BCRP knockdown into the engineered placental barrier model. **e.** Comparison of BCRP expression in the wild-type (WT) and BCRP knockdown (KD) models. Scale bar, 50 μm. The plot shows relative expression of BCRP measured by RT-PCR. **f,g.** Evaluation of β-hCG secretion in the maternal compartment (**f**) and barrier permeability to lucifer yellow (LY) (**g**) after 24 hours of exposure to cadmium. Data were normalized to the untreated control group (0 µM). **h,i.** Comparison of LDH release (**h**) and pro-inflammatory cytokine production (**i**) in the WT and KD models after 24-hour cadmium exposure. Data in (**h**) were normalized to the untreated control group (0 µM). **j,k.** Comparison of endothelial expression of ICAM-1 (**j**) and fibronectin deposition in the stroma (**k**) after 24-hour exposure to 5 µM cadmium. Scale bar, 100 μm. **l.** Demonstration of neutrophil adhesion to the vasculature due to cadmium exposure at 5 µM for 24 hours. Scale bar, 100 μm. **m.** Comparison of cadmium-induced pro-inflammatory cytokine production in the maternal (top row) and fetal (bottom row) compartments in the presence or absence of neutrophils. Data are presented as mean ± SD with *n* = 4. ns = not significant, *\*P < 0.05, **P < 0.01, ***P < 0.001,* and *\*\*\*\* P < 0.0001*.

Motivated by this lack of knowledge, we next delved into how the detected efflux transporters affect responses of the bioengineered placental barrier to cadmium. Based on our PCR data, we hypothesized that during exposure, these proteins are upregulated to increase efflux transport of cadmium from trophoblasts and thus protect the barrier tissue against cadmium toxicity in our model. To test this hypothesis, we individually inhibited the activity of BCRP, MDR1, and MRP, which were selected as three representative efflux transporters, by adding their respective chemical inhibitors in the trophoblast media flowing in the maternal chamber and examined cadmium-induced cellular injury and inflammation in comparison to control without inhibition. In this study, we first treated the model with cadmium chloride at 1 μM, a lower-level exposure condition that did not produce acute tissue injury as evaluated by live/dead staining (**Fig. 2b**) and LDH release (**Fig. 2d**). At this concentration, inhibition of MDR1 and MRP transporters did not change the level of LDH in the effluent of the maternal and fetal compartments (**Fig. 4b**). When the model was modified by using Ko143 to selectively inhibit BCRP function, however, LDH measurements showed nearly two-fold increases on the maternal side, which occurred in the fetal compartment as well although to a lesser extent (**Fig. 4b**). This result demonstrating significantly increased deleterious effects of cadmium in the BCRP-inhibited model was also observed when the cadmium chloride concentration was increased to 5 μM (**Fig. 4b**).

Analysis of cadmium-induced inflammation was performed by measuring pro-inflammatory cytokines in our model exposed to 1 μM of cadmium chloride. Although 5 μM was a threshold concentration necessary to trigger measurable cytokine production in the unmodified control group (**Fig. 3a**), we suspected that the inhibition of efflux transporters may have a significant impact even at lower cadmium levels that normally do not induce inflammatory responses. According to our data, suppressing the activity of MDR1 and MRP failed to alter the cytokine levels in the maternal chamber (**Fig. 4c**). By contrast, exposure to the same concentration of cadmium led to markedly elevated maternal production of all cytokines when BCRP was inhibited (**Fig. 4c**). This condition also resulted in significantly higher levels of all five cytokines in the fetal compartment as well (**Fig. 4c**).

Importantly, these data suggested that BCRP may play a critical role in protecting the maternal-fetal interface from tissue injury and inflammation during cadmium exposure. To validate this, we next used the small interference RNA (siRNA) technique to knock down trophoblast expression of BCRP in our model. Given the difficulty of applying this method to the primary culture of trophoblast cells^54^, we used the BeWo b30 cell line to generate BCRP known-down cells, which were then cultured in the maternal chamber with primary placental fibroblasts and villous endothelial cells in the fetal compartment of the same device (**Fig. 4d**). siRNA did not affect the proliferative capacity and growth rate of these cells in our model, and they formed a confluent monolayer on the membrane surface within 1 day, after which the cells were stimulated with 50 μM forskolin for 3 days to induce syncytialization. The resultant placental syncytium expressed drastically reduced levels of BCRP compared to the control model derived from wildtype BeWo cells (**Fig. 4e**).

The engineered placental barrier was then treated with cadmium chloride at 0.5, 1, 5, and 10 μM. The effect of BCRP knockdown was first assessed by ELISA measurement of β-hCG, which appeared to show decreasing levels of production with the increasing dose of cadmium but without statistical significance, whereas β-hCG in the wild-type model was unaffected at the same levels of exposure (**Fig. 4f**). The threshold cadmium concentration at which the placental barrier began to show compromised integrity was lower in the knock-down model (1 μM) than was measured in the wild-type control (5 μM) (**Fig. 4g**). A loss of BCRP in the trophoblast cells also accentuated cytotoxic effects of cadmium chloride at higher doses (1 and 5 μM) in both maternal and fetal compartments (**Fig. 4h**).

In these higher-level exposure conditions, the knockdown of BCRP resulted in significantly increased production of pro-inflammatory cytokines in the maternal and fetal compartments (**Fig. 4i**). Although the vascularized stroma in the fetal chamber of this model did not contain any modified cells, it exhibited higher levels of cadmium-induced vascular inflammation and matrix deposition when compared to the wildtype control exposed to the same cadmium chloride concentration (**Figs. 4j, 4k**), which is presumably due to elevated cytokine release in the maternal compartment. When the vasculature of this model was perfused with polymorphonuclear neutrophils, we observed large numbers of adherent neutrophils along the endothelial lining (**Fig. 4l**), reminiscent of neutrophil infiltration during fetal inflammatory response reported in pathological studies of the human placenta with acute inflammation^55,56^. The capacity of the recruited neutrophils to exacerbate inflammatory responses to cadmium was demonstrated by the significantly higher levels of pro-inflammatory cytokines in these devices as compared to the control group without neutrophils (**Fig. 4m**).

Taken altogether, these data provide evidence that as one of the most abundant ABC transporters in the human placenta, BCRP plays a central role in protecting the placental barrier from cadmium-induced acute tissue injury and inflammation in our model.

### Metabolomic profiling of the cadmium exposure model – maternal compartment

Traditionally, the primary goal of investigating the adverse effects of toxic metals has been to assess the capacity and potential of the toxicants to induce tissue damage and immune reactions. Increasing evidence shows, however, that toxic metal exposure can also disrupt glucose, lipid, and hormonal homeostasis to elicit aberrant changes in metabolism. In fact, cadmium can cause metabolic dysfunction of the liver by altering the activity of the key enzymes of carbohydrate metabolism^57,58^. Similar effects of cadmium have been demonstrated in other organs involved in metabolic function, such as the intestine and the pancreas^59,60^. Unfortunately, this line of investigation has been scarce in the context of pregnancy, despite the fact that the placenta is a metabolically highly active organ responsible for supporting the growing fetus and itself. Recent epidemiology studies have examined the association of cadmium exposure with relevant conditions such as gestational diabetes^61^, but it remains unknown whether and how cadmium can perturb and exert adverse effects on metabolic activities of the human placenta.

Inspired by this knowledge gap, we conducted unbiased, global analysis of the metabolome of our exposure model, towards the goal of discovering specific markers and molecular signatures indicative of cadmium-induced metabolic changes in the fetal and maternal compartments. In this study, we treated our model established using primary cells with three select concentrations of cadmium chloride (0.1, 1, and 10 μM). For analysis, we used device effluent collected from the maternal and fetal chambers separately over the course of 24-hour exposure (**Fig. 5a**). Dimensionality reduction using principal component analysis (PCA) of our data acquired from the maternal compartment indicated distinct metabolic profiles at different exposure levels, as illustrated by a separation between dose-dependent clusters (**Fig. 5b**). The heatmap representation of 183 most differentially regulated metabolites identified by hierarchical cluster analysis demonstrated that i) the overall patterns of metabolite expression were similar between the untreated control and 0.1 μM groups and that ii) exposure to higher doses (1 and 10 μM) resulted in noticeable changes in their metabolic profiles, especially at 10 μM, in comparison to control and the 0.1 μM group (**Fig. 5c, Supplementary Fig. 4**).

**Figure 5.**
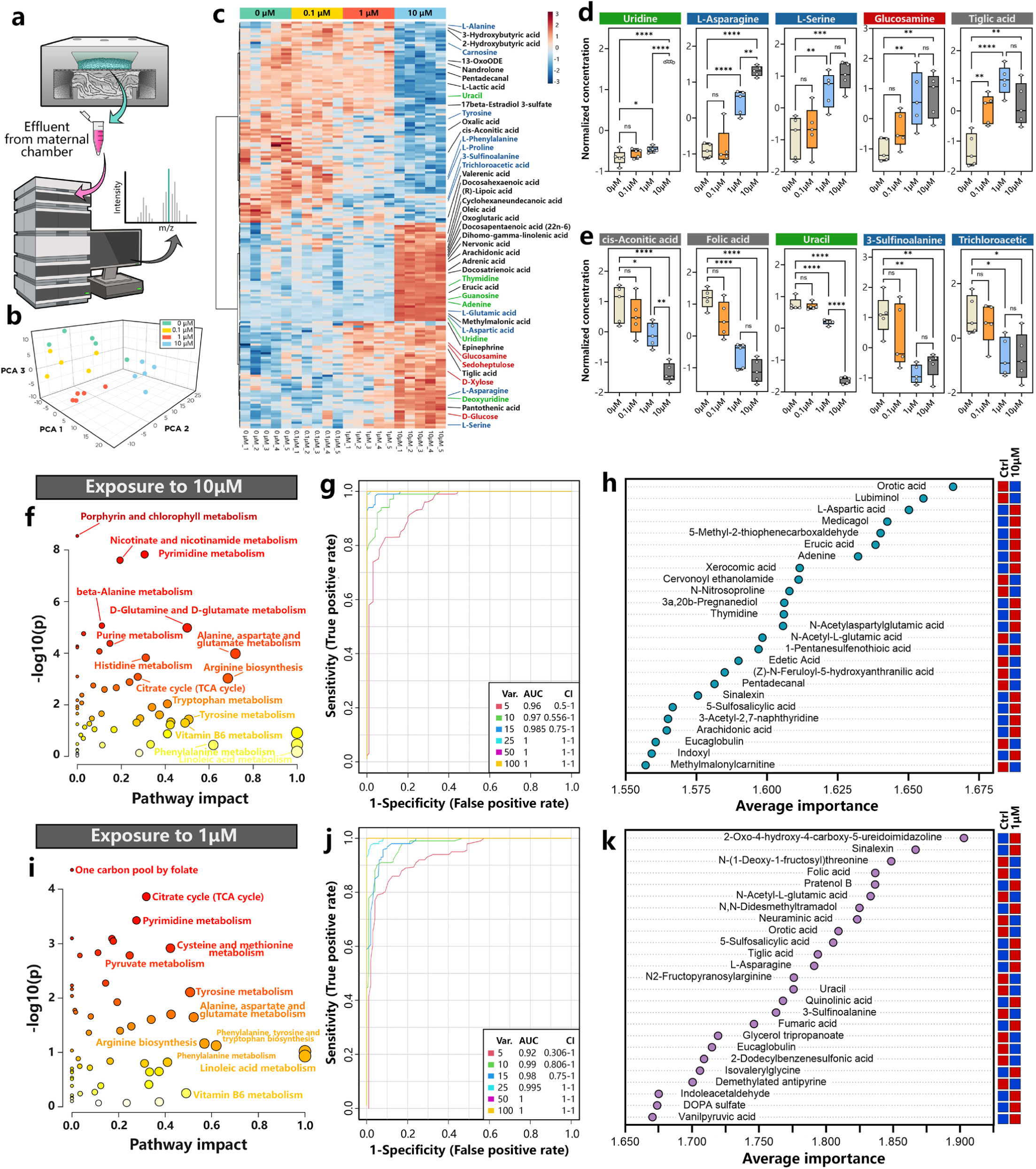
Metabolomic profiling of cadmium-exposed maternal compartment. **a.** Schematic illustration of metabolomic analysis of device effluent collected from the maternal compartment of the engineered placental barrier model. **b,c.** PCA plot (**b**) and heatmap showing relative abundance of top 183 metabolites (**c**) in all conditions tested. **d,e.** Quantification of select metabolite upregulated (**d**) or downregulated (**e**) by cadmium exposure. The color of the box label represents the type of metabolites (blue: peptide metabolites; red: carbohydrate metabolites; grey: lipid metabolites; green: nucleotide metabolites). Data were normalized to the median. **f.** Plot of significantly changed metabolic pathways identified by pathway impact analysis of our model exposed to 10 μM cadmium for 24 hours **g.** ROC curves for biomarker prediction models. Different colors represent models with different numbers of constituent metabolite features. **h.** The list of top metabolites generated by the 25-feature prediction model and ranked based on their predictive accuracy. **i-k.** Pathway impact plot (**i**), ROC curves (**j**), and the list of top metabolite biomarker candidates (**k**) generated by metabolomic analysis of our model exposed to 1 μM cadmium for 24 hours. Box plots show minimum, 25^th^ percentile, mean, 75^th^ percentile, and maximum values. Data are presented as mean ± SD with *n* = 5. ns = not significant, *\*P < 0.05, **P < 0.01, ***P < 0.001,* and *\*\*\*\* P < 0.0001*.

Quantitative comparison of specific metabolites between these groups provided further insight into how cadmium might alter placental metabolism in our model (**Fig. 5d, Supplementary Fig. 5**). For example, our data indicated elevated pyrimidine metabolism as a result of cadmium exposure at 1 and 10 μM, which was evidenced by substantially increased levels of its metabolic product, uridine (**Fig. 5d**). Interestingly, in vivo data from mouse studies have suggested significantly changed pyrimidine metabolism as one of the key metabolic signatures associated with spontaneous abortion due to prenatal exposure to benzophenone-3^62^. Cadmium also increased asparagine and serine metabolism in our model (**Fig. 5d**). Although these pathways have not been investigated in the context of chemical toxicities in the placenta, elevated levels of asparagine and serine were previously shown by metabolomic analysis of the human placenta with fetal growth restriction^63^. Glucosamine was another upregulated marker at higher doses of cadmium known to play a critical role in the synthesis of glycosylated proteins and lipids^64^ (**Fig. 5d**). No evidence exists implicating glucosamine in human placental toxicity of cadmium, but it has been associated with altered fetal growth in mothers with diabetes^65^.

Exposure to the higher doses also led to the reduced activity of several metabolic pathways. This was illustrated by the decreased production of metabolites, such as cis-aconitic acid that plays an essential role in energy production and removal of toxins^66^ and folic acid that is an important contributor to placental development and fetal growth^67,68^ (**Fig. 5e**). In response to lower-level exposure at 0.1 μM, metabolism in the maternal compartment remained largely unaffected, but a small number of metabolites still showed significant changes compared to control, including tiglic acid and D-xylose (**Figs. 5d, Supplementary Fig. 5**). In addition to these examples, our analysis identified many other upregulated or downregulated metabolites with previously unknown links to human placental toxicities of cadmium and other toxic metals (**Supplementary Table 1**).

We then performed pathway impact analysis to gain further insight into the metabolic consequences of cadmium exposure in our model. Comparison of the 10 μM exposure condition with control allowed us to identify and rank 57 metabolic pathways that were most significantly altered by cadmium chloride (**Fig. 5f**). These included pyrimidine and folate metabolism described above, as well as purine metabolism involved in fulfilling the demanding need for energy production in the placenta^69^, nicotinate and nicotinamide metabolism essential for redox reactions that has been shown to change in gestational diabetes^70^, and metabolism of amino acids such as alanine and glutamine that play key roles in supporting fetal growth and metabolism^71^.

Furthermore, we carried out multivariate receiver operating characteristics (ROC) analysis to identify potential metabolite biomarker candidates that may be associated with cadmium toxicity in the human placenta (**Fig. 5g**). This analysis using the 25-feature model showed i) reduction in orotic acid and lubimonol and ii) increased levels of L-aspartic acid, medicagol, and 5-methyl-2-thiophenecarboxaldehyde as the most predictive signatures of metabolic changes due to placental exposure to 10 μM cadmium chloride in our model (**Fig. 5h**).

Finally, the same set of analysis was performed for the intermediate-level (1 μM) exposure condition (**Fig 5i-k**). The metabolic pathways perturbed by this condition included some of those discovered at 10 μM (e.g., pyrimidine and folate metabolism) but data also revealed new ones, including taurine and hypotaurine metabolism, which has been shown by rodent studies to protect trophoblasts from oxidative stress-induced cytotoxicity^74^ and sphingolipid metabolism whose alterations have been associated with gestational diabetes^75^ (**Fig. 5i**). Of note, the top 25 biomarker candidates predicted by ROC analysis for this condition and those for the high-dose condition (10 μM) only shared five metabolites (**Fig. 5k**), suggesting the possibility of developing distinct metabolic signatures that may distinguish placental responses to these two exposure conditions.

### Metabolomic profiling of the cadmium exposure model – fetal compartment

Analysis of metabolites present in the perfusate of the fetal compartment demonstrated similar dose-dependent profiles, but in comparison to what was measured in the maternal chamber, many metabolites began to show noticeable changes even in the lower-level (0.1 μM) exposure condition (**Figs. 6a-6c, Supplementary Fig. 6**). Guanosine was one of them that was significantly upregulated at 0.1 μM, and its level increased further when the concentration was raised to 1 μM (**Fig. 6d**). Cadmium exposure also resulted in substantially increased production of sorbitol at 0.1 μM and higher (**Fig. 6d**). Although these metabolites have not been described in the context of metal toxicity in the placenta, clinical studies have shown their altered production as an indicator of abnormal fetal development and pregnancy complications. Increased levels of guanosine, for example, were among the key findings of metabolomic analysis of human placentas with spontaneous preterm birth^76^ Similarly, sorbitol accumulation in fetal tissues has been suggested to contribute to the development of fetal defects in diabetic pregnancy^77^. In addition to these markers, our analysis showed increased production of many other metabolites without known associations with pregnancy (**Fig. 6d, Supplementary Fig. 7, Supplementary Table 2**).

**Figure 6.**
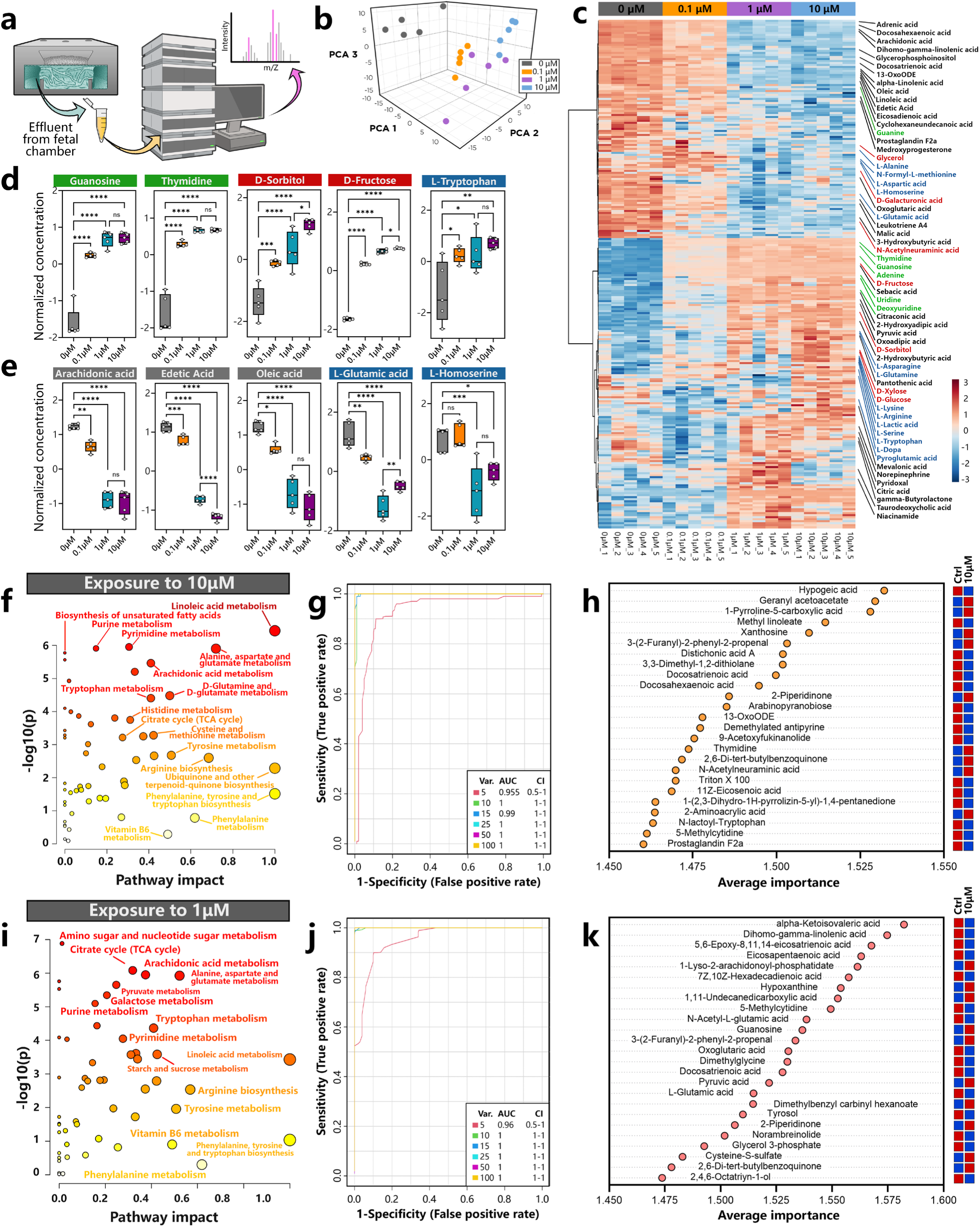
Metabolomic profiling of cadmium-exposed fetal compartment. **a.** Schematic illustration of metabolomic analysis of device effluent collected from the fetal compartment of the engineered placental barrier model. **b,c.** PCA plot (**b**) and heatmap showing relative abundance of top 255 metabolites (**c**) in all conditions tested. **d,e.** Quantification of select metabolite upregulated (**d**) or downregulated (**e**) by cadmium exposure. The color of the box label represents the type of metabolites (blue: peptide metabolites; red: carbohydrate metabolites; grey: lipid metabolites; green: nucleotide metabolites). Data are normalized to the sample median. **f.** Plot of significantly changed metabolic pathways identified by pathway impact analysis of our model exposed to 10 μM cadmium for 24 hours. **g.** ROC curves for biomarker prediction models. Different colors represent models with different numbers of constituent metabolite features. **h.** The list of top metabolites generated by the 25-feature prediction model and ranked based on their predictive accuracy. **i-k.** Pathway impact plot (**i**), ROC curves (**j**), and the list of top metabolite biomarker candidates (**k**) generated by metabolomic analysis of our model exposed to 1 μM cadmium for 24 hours. Box plots show minimum, 25^th^ percentile, mean, 75^th^ percentile, and maximum values. Data are presented as mean ± SD with *n* = 5. ns = not significant, *\*P < 0.05, **P < 0.01, ***P < 0.001,* and *\*\*\*\* P < 0.0001*.

We also found that the levels of several metabolites detected on the fetal side decreased due to cadmium exposure. Examples included: arachidonic acid, an essential fatty acid critical for proper neural development in the fetus^78^, oleic acid that plays an important role in maintaining early-stage pregnancy^79^, glutamic acid that provides a major substrate for energy metabolism in the fetus^80^, and many others that have not been studied in relation to placentation and pregnancy (**Fig. 6e. Supplementary Fig. 7, Supplementary Table 2**). Interestingly, a large fraction of the significant metabolic changes in the fetal compartment were seen in fatty acids, indicating the capacity of cadmium to disrupt lipid metabolism in our model. Consistent with this finding, pathways involved in the metabolism of fatty acids (e.g., arachidonic acid, linoleic acid) were shown by our analysis to be among the most significantly affected pathways in both the 1-μM and 10-μM groups (**Figs. 6f, 6i**). Given that the rest of the highly ranked pathways showed a substantial overlap with those identified in the maternal compartment (**Figs. 5f, 5i**), this result suggests altered fatty acid metabolism as a unique signature of fetal responses to cadmium exposure.

Finally, our biomarker analysis using the ROC curve yielded a set of differentially regulated metabolites associated with fetal-specific effects of cadmium. In case of higher-level exposure at 10 μM, metabolites with the highest predictive values included fatty acids and fatty acid esters, such as hypogeic acid, methyl linoleate, docosatrienoic acid, and docosahexaenoic acid, all of which were present in decreased amounts as a result of cadmium exposure (**Fig. 6h**). The fetal compartment-specific biomarker candidates also included geranyl acetoacetate, 1-pyrroline-5-caboxylic acid, and xanthosine whose levels were elevated by cadmium (**Fig. 6h**). Data from the 1-μM group revealed a different set of fatty acids and other types of metabolites as predictive biomarkers (**Figs. 6j, 6k**). Specifically, fetal metabolic changes due to cadmium exposure at this level were represented by decreased levels of α-ketoisovaleric acid, dihomoγ-linolenic acid, 5,6-epoxy-8,11,14-eicosatrienoic acid, and eicosapentaenoic acid, as well as increased production of 1-lyso-2-arachidonoyl-phosphatidate and hypoxanthine, to name a few with the highest predictive values (**Fig. 6k**). To our knowledge, most of these metabolites and their pathways have not been investigated in the context of pregnancy and fetal development.

## Discussion

Studying physiological or pathophysiological events that occur during human pregnancy is difficult due to ethical restrictions and the limited scope of current clinical investigations, which is fundamental to the widely recognized challenge of basic and translational research in reproductive medicine and related areas^18,81^. This in turn provides a strong justification for developing human cell/tissue-based in vitro models as an alternative approach with unique advantages. Guided by this principle, our work described in this paper demonstrated i) how the salient features of the maternal-fetal interface in the human placenta can be represented and emulated by microengineered cell culture systems and ii) how advanced in vitro capabilities afforded by such systems may be leveraged to model and probe placental toxicity of environmental metals. Specifically, we created a specialized in vitro model of cadmium exposure in the human placenta and used this system in conjunction with various analytical techniques to simulate and understand its adverse effects at the tissue, cellular, and molecular levels.

Given that a few studies have previously demonstrated the ability to model the human placental barrier in vitro using Transwell inserts or microdevices^82,83^, it may be worthwhile to discuss how our model presented here compares to those existing systems. Perhaps, the most fundamental difference lies in the fact that our model was constructed using primary human placental cells from chorionic villi and grown under more physiologically relevant culture conditions, namely at lower (5%) oxygen tension. This is in contrast to other published studies that commonly used human trophoblast cell lines, such BeWo and JEG, and macrovascular placental endothelial cells cultured at 20% oxygen^84–87^. As shown in our work, the physiological oxygen environment allowed primary trophoblasts to retain their proliferative capacity and also made it possible for our model to more faithfully recapitulate physiological properties of the placental barrier in vivo, including spontaneous syncytialization and higher β-hCG production in primary cells.

The only exception was our investigation of the role of BCRP in which we established our models using the BeWo cell line. Use of these cells was necessary due to the well-known technical difficulty of knocking down the expression of the efflux transporter in primary trophoblasts^54^. Given the focus of our work on cadmium toxicity, however, we recognize that work remains to be done to systematically examine how responses of our primary cell-based models to cadmium compare to those in cell line-based models. While further investigation is necessary, our data showed higher levels of pro-inflammatory cytokine production in the BeWo cell line-based models compared to primary culture-based devices exposed to the same concentrations of cadmium (**Supplementary Fig. 8**).

The vascularized stroma is another unique feature of our model not demonstrated in previous studies. This element allowed us to mimic the structural organization and 3D tissue-specific microenvironment of the maternal-fetal interface in the human placenta with higher fidelity, which may be considered a notable improvement and difference from the 2D culture configuration commonly used in existing models in which two opposing monolayers of trophoblast cells and endothelial cells are formed on either side of a semipermeable membrane ^26,82,85–88^. On the other hand, this comparison begs the question of whether and how this more realistic architecture of the model affects its biological activities, especially its responses to cadmium examined in our study. As can be seen in **Figs. 2b-d**, the stromal compartment was in fact found to influence responses of the in vitro placental barrier to maternal cadmium by making the tissue more resistant to cytotoxic effects of cadmium as compared to the 2D models created using conventional platforms, such as Transwell. Considering that substantial tissue injury and inflammation due to cadmium exposure at relatively low levels (e.g., 1 μM) observed in the Transwell model has not been reported in human placentas, this may be interpreted as the capacity of our engineered model to produce more in vivo-like responses.

The stromal tissue in our system also provided a means to incorporate other relevant cell types and examine their contributions to placental responses to cadmium, which was demonstrated by our study of Hofbauer cells (**Fig. 3e**). This configuration also made it possible to measure other adverse reactions beyond cytotoxicity and inflammation, as shown by the demonstration of cadmium-induced matrix remodeling (**Fig. 3k**), which has been described in animal models^88,89^. It is suspected that these advanced features may serve to make our model and its produced results more representative of what happens in the native tissue, but this should be validated through comparative analysis of our findings to those from cadmium-exposed human placentas. Unfortunately, these types of studies are currently challenging due to a scarcity of such human data, which represents an important issue that needs to be addressed in future studies.

From a conceptual standpoint, we believe that our work embodies an important advance in that it provides a rare example of leveraging recent developments in microphysiological systems technology and related in vitro approaches towards advancing our ability to address major questions in the study of environmental impact on human pregnancy. The ability to engineer more representative, realistic models of the human reproductive system would have a significant impact on this line of investigation, but the utility of such advanced models has focused mainly on disease modeling and preclinical drug testing. In this regard, our work may represent a promising yet underexplored area of research efforts for both model developers and environmental health scientists. Based on the proof-of-concept demonstration presented here, greater attention should be paid to the possibility of constructing more advanced or different types of human reproductive tissue models and harnessing their potential as uniquely powerful platforms to investigate the unknowns of how existing and emerging environmental materials may affect human reproduction.

To discuss specific findings from the exposure model, our experiments conducted using a range of cadmium concentrations verified the general expectation that substantially higher levels of maternal cadmium, such as 10 or 50 μM tested in our study, may have the capacity to induce acute cellular injury in the human placental barrier. Given that human environmental exposure to cadmium usually occurs at lower levels (< 10 μM)^35^, however, the prevailing thought is that that cadmium may generate placental toxicity by disrupting biological signaling that plays an important role in regulating the phenotype and function of the placental barrier, rather than directly harming the tissue^90^. Supporting this hypothesis, our data showed that lower-level exposure conditions can trigger non-injurious yet adverse cellular and tissue responses, such as reduced hormone production, decreased maternal-to-fetal glucose transfer, increased cytokine production, and altered expression of trophoblast membrane transporters, without showing any significant changes in cell viability. Although not in the context of cadmium exposure, such maladaptation of the human placenta has been suggested to play a causative role in adverse outcomes of human pregnancy^91^. Based on these findings, we can suspect that altered placental activities due to cadmium can irreversibly dysregulate placental function over longer periods of time, which may contribute to the development of pregnancy complications associated with cadmium exposure. The goal of our study was to simulate acute exposure, but significant value can be added to our current data if the system could be modified to prolong the duration of culture/cadmium treatment and examine longer-term responses of the engineered human placental barrier.

Also important to discuss in this context are our metabolomics data. The results of this analysis provided further evidence of cadmium toxicity by demonstrating significantly altered metabolic activities of the placental barrier. Research has shown adverse effects of cadmium and other environmental metals on metabolic function of various organs^92^, but to our knowledge, its capacity to induce abnormal metabolic changes in the human placenta during pregnancy has not been described in the literature. What is more important to highlight is a high degree of similarity between some of the specific changes observed in our model and those reported in metabolomic interrogation of human placentas with pregnancy complications^93^. As can be seen in many examples described in the Results section, several of the differentially regulated metabolites due to cadmium exposure in our model have previously been identified as signatures of aberrant metabolic changes associated with spontaneous preterm birth, gestational diabetes, spontaneous abortion, or fetal growth restriction. Although our system is unable to model direct effects of cadmium on fetal development, these findings suggest disrupted placental metabolism as an important consideration for understanding the biological basis of how cadmium exposure can lead to adverse pregnancy outcomes, which remains a major knowledge gap in the field.

On a related note, cadmium-induced disruption of lipid metabolism in the fetal compartment was one of the key findings of our analysis. This result bears significance in that it provides additional new insight into how prenatal cadmium exposure may have adverse effects on fetal development. The placenta is known to take up lipids from the maternal circulation and convert them into free fatty acids, which are then used to support its own growth and transported to the fetal circulation to meet the high metabolic needs of the growing fetus^94,95^. This highly regulated process of lipid metabolism and transport in the placenta becomes even more important at late stages of gestation as lipids play a vital role in proper neurodevelopment in the fetus^95,96^. Studies have shown that placental processing of lipids can be dysregulated by maternal conditions such as obesity and that this is important for understanding placental maladaptation and pregnancy complications common in such conditions^97^. Whether and how placental lipid metabolism is affected by prenatal metal exposure, however, is currently unknown. By revealing significantly changed levels of fatty acids in the fetal compartment, our data provide in vitro evidence suggesting that cadmium may have the capacity to disrupt lipid metabolism in the human placenta and that this may help us better understand the established link between cadmium exposure and adverse pregnancy outcomes.

Investigation of placental ABC transporters was another important goal of our study. Although extensive work has been done to study the role of these transmembrane proteins in regulated cellular export of ions, macromolecules, toxins, and drugs in various organs^98^, there is a lack of evidence about their capacity to mediate cadmium efflux from the human placental barrier. In particular, BCRP is one of the two most abundant ABC transporters expressed in the human placenta that has recently been shown to protect trophoblasts from cadmium^99^, but much remains to be learned about its activity in the integrated context of the human placental barrier exposed to maternal cadmium. Our study provides in vitro data that may be useful for addressing this knowledge gap. We showed that BCRP serves to reduce the extent of cadmium toxicity in both the maternal and fetal compartments of the engineered placental barrier during low-dose exposures. We also found that BCRP plays a more important protective role than some of the other ABC transporters expressed in our system, such as MDR1 and MRP2. In vivo translatability of these findings remains to be further investigated, but we believe that they still provide scientific value by reinforcing the general notion of ABC transporters as an inherent protective mechanism against foreign substances while also offering context-specific new insight into the importance of BCRP-mediated efflux function in understanding placental toxicity of cadmium.

For further development of this study, it would be interesting to delve into the detailed mechanisms of BCRP-mediated efflux transport of cadmium in the trophoblast layer of our model towards the goal of exploring the feasibility of modulating this process to increase the protective capacity of BCRP against cadmium toxicity. In case of MRP1 and ABCB6, both of which belong to the same ABC transporter superfamily as BCRP, researchers have discovered in other organs that cadmium efflux by these transporters only occurs when cadmium binds to its physiological substrate, glutathione (GSH), and forms a complex^100^. Based on this finding, there has been discussion about the possibility of developing potential strategies for cadmium detoxification that use externally administered substances to promote GSH production^101–103^. The same approach may be extended to cadmium efflux mediated by BCRP in the human placenta, but this will first require in-depth investigation of this transport process to identify its endogenous substrates that can be modulated externally.

Expanding the scope of our metabolomics work may be another promising area of investigation for future studies. This study yielded a set of dose-specific metabolite biomarkers indicative of cadmium-induced metabolic changes in the maternal and fetal compartments of our model. Given that the maternal-fetal interface undergoes substantial structural and functional changes over the course of pregnancy, it would be interesting to modify the design of our current system and simulate cadmium exposure at different stages of human pregnancy, which could then be analyzed using the same techniques to produce multiple distinct sets of stage-specific metabolite biomarkers. For further enrichment of data, efforts could also be made in future studies to vary the duration of cadmium exposure in our models to measure how metabolic profiles of the fetal and maternal compartments change over time. Although already known to be challenging, a major step forward for this type of work would be in vivo validation of these biomarker candidates in human subjects. If possible, such studies will generate pre-clinical data that may play an instrumental role in developing clinical biomarkers for the assessment and prediction of human placental toxicity of cadmium, which currently do not exist. They may also provide opportunities to assess the potential of our and other similar emerging models of human reproductive tissues as biomarker discovery platforms.

On a final note, we point out that by design, our model is a simplified representation of remarkably complex placental anatomy and biology and is therefore limited in its ability to reproduce the reality of what happens in the native human placenta during cadmium exposure. As discussed above, we recognize several limitations in our current system and study, including, for example, no consideration of dynamic evolution of the maternal-fetal interface during the progression of pregnancy, short durations of cadmium exposure, a lack of attention to the effect of hemodynamic forces, inability to model short- and long-term adverse effects of cadmium on fetal development, and so on. We think that the design flexibility and controllability of the 3D culture system in our model will make it feasible to address most of these issues, with the exception of modeling fetal responses to cadmium. What features and capabilities to include or exclude in future studies, however, should be determined by the specific questions at hand.

In closing, we believe that further development of this work through concerted collaborative efforts between engineers, clinicians, and toxicologists will provide opportunities to create powerful, innovative research tools that can change and advance our ability to model, probe, delineate, and modulate the biology of how our dynamically changing environment impacts human reproduction.

## Materials and methods

### Fabrication of cell culture devices

Devices used for culturing human placental cells in this study were fabricated using soft lithography. Briefly, poly(dimethyl siloxane) (PDMS, Sylgard 184, Dow Corning) monomer base was mixed with a curing agent (10:1, w/w) and poured onto 3D-printed molds manufactured by stereolithography (Protolabs). After degassing in a desiccation chamber (Bel-Art Inc.) for 1 hour, PDMS was cured in an oven at 65 °C for 2 hours. Subsequently, a Whatman Cyclopore Polycarbonate Thin Clear membrane with 1 µm pores (Cytiva, USA) was punched using a 7mm biopsy punch (Acuderm Inc., USA) and sealed against the middle PDMS layer using uncured PDMS as a glue. Next, fluidic access ports were generated in the topmost PDMS layer (tubing layer) using a 1 mm biopsy punch (Integra, Inc.), and this layer was bonded to the upper PDMS layer containing the maternal chamber. The device assembly was then placed in an oven for 1 hour and stored in a container until use.

### Cell culture

Human primary cells used in this study were obtained from the Amnion Foundation (NC, USA) that included human cytotrophoblast cells (Cat #1230), placental stromal fibroblasts (Cat# 1250), MVECPRO2, germinal-origin microvascular endothelial cells (Cat #1245), and Hofbauer cells (Cat #1220). Human placental microvascular endothelial cells and placental fibroblasts were seeded and grown in Corning TC-treated T-75 tissue culture flasks. The endothelial cells and fibroblasts were cultured in microvascular endothelial cell growth medium (MVECPRO2: Lonza EGM-2 MV) and fibroblast media (Stromal cells/fibroblasts; Lonza FGM-2), respectively. These cells were used for creating engineered placental tissues after one passage. Human cytotrophoblast cells were cultured using cytotrophoblast culture media (CTBPRO2GRO: Amnion Foundation) in an incubator maintained at 5% oxygen.

### Tissue production in the fetal chamber

After device fabrication, the PDMS chambers were filled with 70% ethanol and incubated for 1 minute, after which ethanol was removed by applying vacuum aspiration to access ports. Subsequently, the device was sterilized for 20 minutes using ultraviolet (UV) light. To generate vascularized stromal tissues in the fetal compartment, a 100-µl mixture of 10 mg/ml fibrinogen (Millipore Sigma Cat. F8630-5G) in DPBS, thrombin (10 U/ml), microvascular endothelial cells (5 × 10^6^ cells/ml), and placental fibroblasts (5 × 10^6^ cells/ml) were injected into the fetal chamber of the lower device layer compartment of a microfabricated device. For devices containing Hofbauer cells, primary human Hofbauer cells (Amnion Foundation) were first labeled with CellTracker™ red CMFDA and added to the ECM-cell mixture at a final concentration of 1 × 10^6^ cells/ml, which was then injected into the fetal chamber of our device. The seeded device was placed in an incubator at 37 °C for 15 minutes to induce gelation of the mixture. Subsequently, the side channels were seeded with endothelial cells (5 × 10^6^ cells/ml). After cell attachment, the fetal chamber of the device was connected to a computer-controlled flow pump driven at a volumetric flow rate of 100 μl/h.

### Transwell culture

Transwell culture of primary human placental cells was established for comparative analysis of biological responses to cadmium. This model was created in a Transwell insert (Corning, NY, USA catalog #38024) containing a semipermeable membrane with a pore size of 0.4 μm. First, microvascular placental endothelial cells were plated on the lower side of the membrane insert at a concentration of 5 × 10^6^ cells/ml while the insert was kept inverted. After 1.5 hours of incubation at 37 °C, 5% CO2 to permit cell attachment, the endothelial cell-seeded insert was placed in a culture well containing fresh EGM-2MV media. Following this step, trophoblast cells were seeded onto the upper side of the membrane at 5 × 10^6^ cells/ml. During culture, media were changed every 24 hours until 100% confluency was reached.

### Cadmium exposure

The cadmium used in this study was sourced from CdCl2 (Sigma-Aldrich, catalog #202908), prepared as a stock solution in distilled deionized water, and appropriately diluted with medium before application. To generate an exposure model, the maternal chamber was perfused with trophoblast media containing defined concentrations of cadmium chloride (0, 0.1, 0.5, 1, 5, 10, and 50 μM). During exposure, the devices were maintained at 37 °C and 5% CO2.

### Immunofluorescence staining and microscopy

For immunofluorescence staining, tissues were first fixed in a 4% PFA solution at 4 °C for 4 hours and then permeabilized in 1% Tween-20 and 10% BSA at 4°C for 2 hours. Primary antibodies were applied at manufacturer-recommended dilutions or at 1:100 dilution, whichever was more concentrated, and incubated at 4 °C overnight. On the next day, tissues were washed with DPBS containing 0.1% Tween-20 and 10% BSA and left overnight. Secondary antibodies, if required, were incubated overnight, followed by another day of washing in the same solution. DAPI and Phalloidin were added at dilutions of 1:5000 and 1:1000, respectively. Following staining, imaging was conducted using confocal microscopy (Zeiss LSM 800) with 10X 0.45 NA and 63X 1.4 NA objectives.

Antibodies used for immunofluorescence analysis included CD31 (Alexa Flour 488 Anti-CD31, Abcam ab215911), E-Cadherin (Abcam ab40772 and ab1416), GLUT-1 (Abcam ab115730), CD163 (Abcam ab182422, Invitrogen MA5-17716), ZO-1 (Invitrogen 33-9100, Invitrogen 61-7300), ICAM-1 (Invitrogen 1A29), Fibronectin (Invitrogen 53-9869-82), α-SMA (Invitrogen 14-9760-82, R&D Systems MAB1420), and BCRP (Santa Cruz sc-18841). The complete list of antibodies is shown in **Supplementary Table 3**.

### Measurement of cell viability, LDH release, and apoptosis

The viability of trophoblast cells in the maternal compartment was analyzed using the Live/Dead Cell Viability Kit (Life Technologies) following the manufacturer’s instructions. Briefly, the cells in the device were washed with PBS and stained with PBS containing 4 μM ethidium homodimer and 2 μM calcein AM at 37 °C for 30 min. Subsequently, the cells were rinsed with fresh PBS and imaged using a laser scanning confocal microscope (LSM 800, Carl Zeiss, Jena, Germany). To measure LDH release from the engineered placental tissues, perfusate from the device was collected from the outlet access ports and analyzed by the Cytotoxicity Detection KitPLUS assay (LDH, Roche, 04744926001) using the manufacturer-provided protocol. Analysis of caspase-3/7 was performed by using the Caspase-Glo^®^ 3/7 Assay (Promega). An equal volume of reagent was added and gently mixed at 37 °C for 2 hours according to the manufacture’s protocol. Luminescence generated by apoptotic cells was detected and quantified using a fluorescence plate reader (Infinite M PLEX, Tecan, Männedorf, Switzerland)

### Analysis of secretory products

To measure β-hCG production by trophoblast cells in our model, effluent from the maternal chamber was collected on day 10 of culture. The collected media samples were then analyzed by the human β-hCG ELISA Kit (DKO014, DiaMetra) following the manufacturer-provided protocol to measure the concentration of β-hCG. For analysis of pro-inflammatory cytokines, device effluent collected from the maternal and fetal chambers was measured using the following ELISA kits: Human IL-1β ELISA Kit (RAB0273, Sigma Aldrich), Human IL-6 ELISA Kit (RAB0306, Sigma Aldrich), Human IL-8 ELISA Kit (RAB0319, Sigma Aldrich), Human TNF Alpha ELISA Kit (ab181421, abcam), and Human TGF Beta 1 ELISA Kit (DY240, R&D Systems).

### Measurement of barrier function

The structural integrity of the engineered placental barrier was assessed by measuring electrical resistance between the maternal and fetal chambers. Briefly, Ag/AgCl electrodes (0.008’’ diameter, A-M Systems, WA, USA) connected to a multimeter (Fluke, USA) were inserted into the access ports of the maternal and fetal chambers. Resistance evaluated from an empty device without any cells was subtracted from each measurement to calculate net resistance, which was then multiplied by the surface area of the culture chamber to compute the final values expressed in Ω × cm^2^.

Barrier permeability was evaluated by measuring transfer of tracer dyes including Lucifer Yellow CH (Invitrogen, L453) and Bovine Serum Albumin 594 conjugated (BSA, Biotium, cat # 20290) from the maternal to fetal compartments. Briefly, a solution containing 1 mM Lucifer Yellow in HBSS/HEPES was introduced into the maternal chamber, while the fetal chamber was filled with HBSS/HEPES. Over a period of 2 hours, outflow from the fetal chamber was collected, and its fluorescent intensity was measured using a microplate reader (Tecan, excitation 485 nm, emission 530 nm).

### RT-PCR analysis

For RNA isolation from the maternal compartment, trophoblast cells were treated with trypsin at 37°C for 5-10 minutes and then harvested from the device. Total RNA for PCR analysis was extracted using the High Pure RNA Isolation Kit (Roche) following the manufacturer’s instructions. In the fetal compartment, the fibrin gel was dissolved with an enzyme solution mixed with trypsin. Subsequently, total RNA was isolated using the RNeasy Plant Mini Kit (Qiagen) according to the manufacturer’s protocol. The RNA was then converted to cDNA using the Transcriptor First Strand cDNA Synthesis Kit (Roche). Quantitative PCR reactions were set up with the FastStart Universal SYBR Green Master (Roche) using the primers listed in **Supplementary Table 4**. The relative expression level of each gene was calculated using the comparative threshold cycle (ΔCt) method using GAPDH as a housekeeping gene.

### Analysis of glucose transport

To measure maternal-to-fetal transfer of glucose in our model, the maternal chamber was perfused with trophoblast media supplemented with D-glucose (Gibco) to generate a final glucose concentration of 450 mg/dl. Culture media flowing through the fetal compartment contained 100 mg/dl of glucose. Over a 2-hour period, outflow from both the maternal and fetal chambers was collected, and its glucose concentration was measured using a digital glucose meter (Auvon, USA). The rate of maternal-to-fetal glucose transfer was evaluated by calculating the percent increase in fetal glucose concentration over the perfusion period.

### Generation of BCRP knock-down model

The human choriocarcinoma BeWo cells were cultured in DMEM/F12-K (Life Technologies) supplemented with 10% fetal bovine serum and 1% penicillin-streptomycin in an incubator at 5% CO2 and 37°C. BeWo knock-down cells were generated using BCRP lentiviral shRNA particles (sc-41151-V, Santa Cruz Biotechnology, Dallas, TX) as previously described^104^. For device seeding, the sterilized membrane surface in the maternal chamber was coated with gelatin (0.1% Gelatin Solution) and fibronectin (20 μg/ml) for 2 hours and seeded with BeWo cells at 2.5 x 10^6^ cells/ml, after which the device was incubated at 37 °C for at least 1.5 hours to allow the cells to adhere to the membrane surface. Once cell attachment was confirmed, the chamber was connected to a pump for continuous perfusion at a flow rate of 100 μl/h. To induce syncytialization, the BeWo cells in the maternal chamber were perfused with DMEM/F12-K medium containing 50 x10^−6^ M forskolin (Sigma).

### Plasmonic analysis of cadmium

Gold nanoparticles (AuNPs) with a diameter of 44 nm were synthesized using a standard sodium citrate reduction method^105^. The AuNPs were modified with poly(ethylene glycol) (PEG) following a previously described protocol with some modifications^106^. Briefly, 5 ml of the as-prepared AuNPs solution was stirred vigorously at room temperature. Next, 50 μl of poly(ethylene glycol) 2-mercaptoethyl ether acetic acid (Mn 2,100, Sigma Aldrich) was added dropwise, and the solution was stirred overnight. The solution was then sonicated for 3 minutes and centrifuged at 2,800 rcf for 10 minutes, twice. Subsequently, 700 μl of the PEG-modified AuNP solution was mixed with 300 μl of the medium and left for 3 hours prior to absorbance measurement. The absorbance spectra were measured using either a Cary 60 UV-vis spectrophotometer or a Cary 5000 UV-vis spectrophotometer (Agilent Technologies Inc.).

### Introduction of human peripheral blood neutrophils

Human peripheral blood neutrophils were isolated from the whole blood provided by a healthy donor using a the EasySep™ Direct Human Neutrophil Isolation Kit (StemCell Technologies, Cat# 19257). To introduce the neutrophils into our exposure model, we stained the cells with Hoechst and suspended them in EGM-2 medium at the final concentration of 2.5-3 × 10^6^ cells/ml. The cells were then injected into the vessels through the inlet access port for one of the sides microchannels and allowed to flow through the vasculature for 24 hours while the device was maintained in a cell culture incubator. At the completion of perfusion, the device was washed with DPBS three times and imaged to quantify the number of adherent neutrophils.

### Supernatant metabolite extraction for metabolomics analysis

Samples of culture media were collected from individual microdevices at defined time points and stored at -80°C. 5 μl of media was added to 120 μl of −20°C 25:25:10 (v/v/v) acetonitrile:methanol:water solution, vortexed for 10 seconds, and put on ice for at least 5 minutes. The resulting extract was centrifuged at 16,000 × g for 20 minutes at 4°C, and the supernatant was transferred to tubes for LC-MS analysis. A procedure blank sample was generated identically without culture media, which was used later to remove the background ions.

### Metabolite measurement by LC-MS

Metabolites were analyzed using a Vanquish Horizon UHPLC System (Thermo Scientific) coupled to an Orbitrap Exploris 480 Mass Spectrometer (Thermo Scientific). Waters XBridge BEH Amide XP Column (particle size, 2.5 μm; 150 mm (length) × 2.1 mm (i.d.)) was used for hydrophilic interaction chromatography (HILIC) separation. Column temperature was kept at 25 °C. Mobile phases A = 20 mM ammonium acetate and 22.5 mM ammonium hydroxide in 95:5 (v/v) water:acetonitrile (pH 9.45) and B = 100% acetonitrile were used for both ESI positive and negative modes. The linear gradient eluted from 90% B (0.0–2.0 min), 90% B to 75% B (2.0– 3.0 min), 75% B (3.0–7.0 min), 75% B to 70% B (7.0–8.0 min), 70% B (8.0–9.0 min), 70% B to 50% B (9.0–10.0 min), 50% B (10.0–12.0 min), 50% B to 25% B (12.0–13.0 min), 25% B (13.0–14.0 min), 25% B to 0.5% B (14.0–16.0 min), 0.5% B (16.0–20.5 min), then stayed at 90% B for 4.5 min. The flow rate was 0.15 mL/min. The sample injection volume was 5 μL. ESI source parameters were set as follows: spray voltage, 3200 V or −2800 V, in positive or negative modes, respectively; sheath gas, 35 arb; aux gas, 10 arb; sweep gas, 0.5 arb; ion transfer tube temperature, 300 °C; vaporizer temperature, 35 °C. LC–MS data acquisition was operated under full scan polarity switching mode for all samples. The full scan was set as: orbitrap resolution, 120,000 at m/z 200; AGC target, 1e7; maximum injection time, 200 ms; scan range, 60– 1000 m/z.

### Analysis of metabolomics data

LC-MS raw data files (.raw) were converted to mzXML format using ProteoWizard (version 3.0.20315)^107^. El-MAVEN (version 0.12.0)^108^ was used to generate a peak table containing m/z, retention time, and intensity for the peaks. Parameters for peak picking were the defaults except for the following: mass domain resolution, 5 ppm; time domain resolution, 10 scans; minimum intensity, 10,000; and minimum peak width, 5 scans. The resulting peak table was exported as a .csv file. Peak annotation of untargeted metabolomics data was performed using NetID^108^ with default parameters. Statistical analyses were performed using MetaboAnalyst 5.0^109^.

### Statistical analysis and reproducibility

A minimum of three biological replicates were used for each experimental group. All data in the paper are represented as mean ± standard deviation (SD). Data were analyzed with Student’s t-test and with one-way and two-way ANOVA followed by Tukey’s post-hoc test for multigroup pairwise comparisons. Differences were considered statistically significant at p-values < 0.05. *P ≤ 0.05, **P ≤ 0.01, ***P ≤ 0.001, ****P ≤ 0.0001. GraphPad Prism (ver. 9; GraphPad Software) was used for statistical analysis.

## Acknowledgements

This work was supported by the National Institutes of Health (NIH) (1R01ES029275, 1UC2HD113039, P30ES013508, P30ES005022, UM1TR004789) (DDH, LMA, PF), the National Science Foundation (CMMI:15-48571) (DDH), the Ministry of Trade, Industry & Energy of the Republic of Korea (DDH, TK), and the University of Pennsylvania. P.F. is the recipient of the CEET Mentored Scientist Transition Award (MSTA) from the Center of Excellence in Environmental Toxicology at the University of Pennsylvania.

## Author contributions

P.F. designed the research, performed the experiments, analyzed the data, and wrote the manuscript. M.Y. performed the experiments and analyzed the data. W.D.L. and J.D.R. conducted the metabolomics experiments and analyzed the data. K.W. and T.K. carried out the plasmonic detection experiments and analyzed the data. L.M.A. provided input into the design of research and analysis of data. D.D.H. designed the research, analyzed the data, and wrote the manuscript.

## Data availability

The datasets generated and/or analyzed during the current study are available from the corresponding author upon reasonable request.

