## Supplemental Information for "A bioengineered model of human placental exposure to environmental metals during pregnancy"

**Supplementary Information includes:**

**Supplementary Figures 1 to 8:**
**Supplementary Figure 1.** Injection and spatial confinement of cell-ECM mixture
**Supplementary Figure 2.** Effect of flow culture conditions
**Supplementary Figure 3.** Plasmonic detection and quantification of cadmium
**Supplementary Figure 4.** Heatmap of metabolites in the maternal compartment
**Supplementary Figure 5.** Comparison of select metabolites in the maternal compartment
**Supplementary Figure 6.** Heatmap of metabolites in the fetal compartment
**Supplementary Figure 7.** Comparison of select metabolites in the fetal compartment
**Supplementary Figure 8.** Comparison of cadmium-induced cytokine production in BeWo cell- and primary cell-based models

**Supplementary Table 1 to 4:**
**Supplementary Table 1.** Select maternal metabolites and their association with human placental toxicity of environmental metals
**Supplementary Table 2.** Select fetal metabolites and their association with human placental toxicity of environmental metals
**Supplementary Table 3.** Key resources
**Supplementary Table 4.** List of primers

### Supplementary Figure 1. Injection and spatial confinement of cell-ECM mixture

Time-series images demonstrating the injection of cell-ECM mixture solution into the fetal chamber of the device. Injection only occurs in the bottom layer separated from the top compartment by a permeable membrane – the top and membrane layers are not clearly visible in these images. Scale bars, 1 mm.

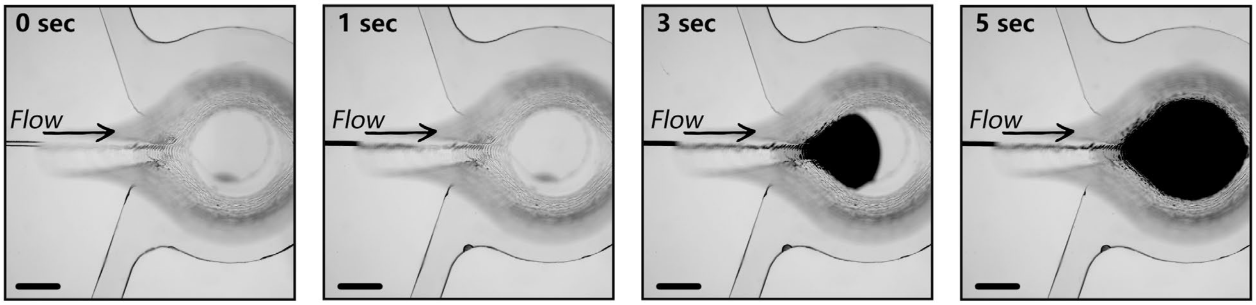


### Supplementary Figure 2. Effect of flow culture conditions

**a.** Representative confocal images comparing vascular formation under static and dynamic conditions on day 7 (left) and quantification of select vascular features (right). Green fluorescence in the confocal images shows immunostaining of CD31 expressed by endothelial cells (scale bar = 100μm). **b.** Comparison of microvilli formation on the apical surface of trophoblasts under static and flow culture conditions. Microvilli in the confocal images are shown by immunofluorescence staining of actin. Blue shows nuclear staining. The micrograph showing a top view of microvilli in the dynamic culture condition was repeated from **Fig. 1I**. Data are presented as mean ± SD with $n = 4$. $^{**}P < 0.01$, and $^{****}P < 0.0001$.

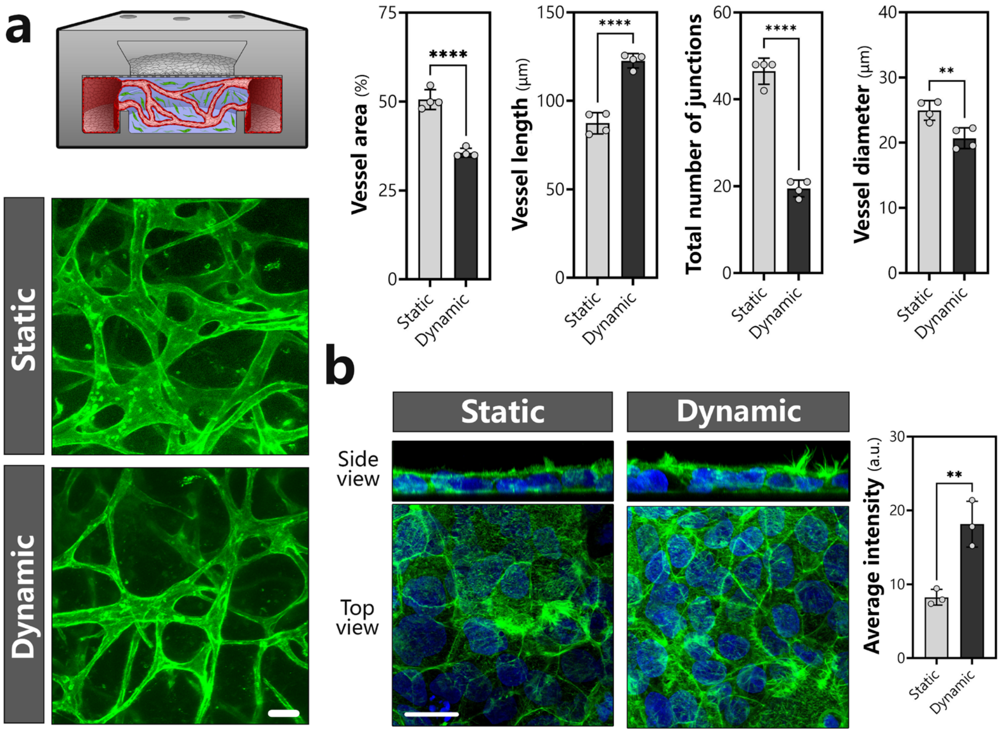

### Supplementary Figure 3. Plasmonic detection and quantification of cadmium

**a.** Gold nanoparticles (AuNPs) functionalized with carboxylic acid-terminated poly(ethylene glycol) (PEG) surface groups were generated using a sodium citrate reduction method. In the presence of cadmium, the distance between these particles decreases due to the chelation of cadmium ions by the carboxylate terminals of PEG, providing a basis for plasmonic detection of cadmium in our effluent samples. **b.** Absorbance spectra measured after mixing AuNP-containing culture media with different concentrations of cadmium ions. As the concentration of cadmium increases, the intensity of the surface plasmon resonance (SPR) band at 535 nm ($A_{max}$) decreases, whereas SPR intensity at 800 nm ($A_{800}$) increases. **c.** During 24-hour cadmium exposure in our model, device effluent was collected from the maternal and fetal chambers of our device and mixed with AuNP solution. Subsequently, $A_{800}$ and $A_{max}$ were measured for both compartments, and their ratios (shown as M and F in the plot) were used in conjunction with the known concentration of cadmium in the maternal chamber ($Cd_{maternal}$) to calculate the cadmium concentration in the fetal chamber ($Cd_{fetal}$), which also allowed us to estimate the rate of maternal-to-fetal cadmium transfer shown in **Fig. 2f**.

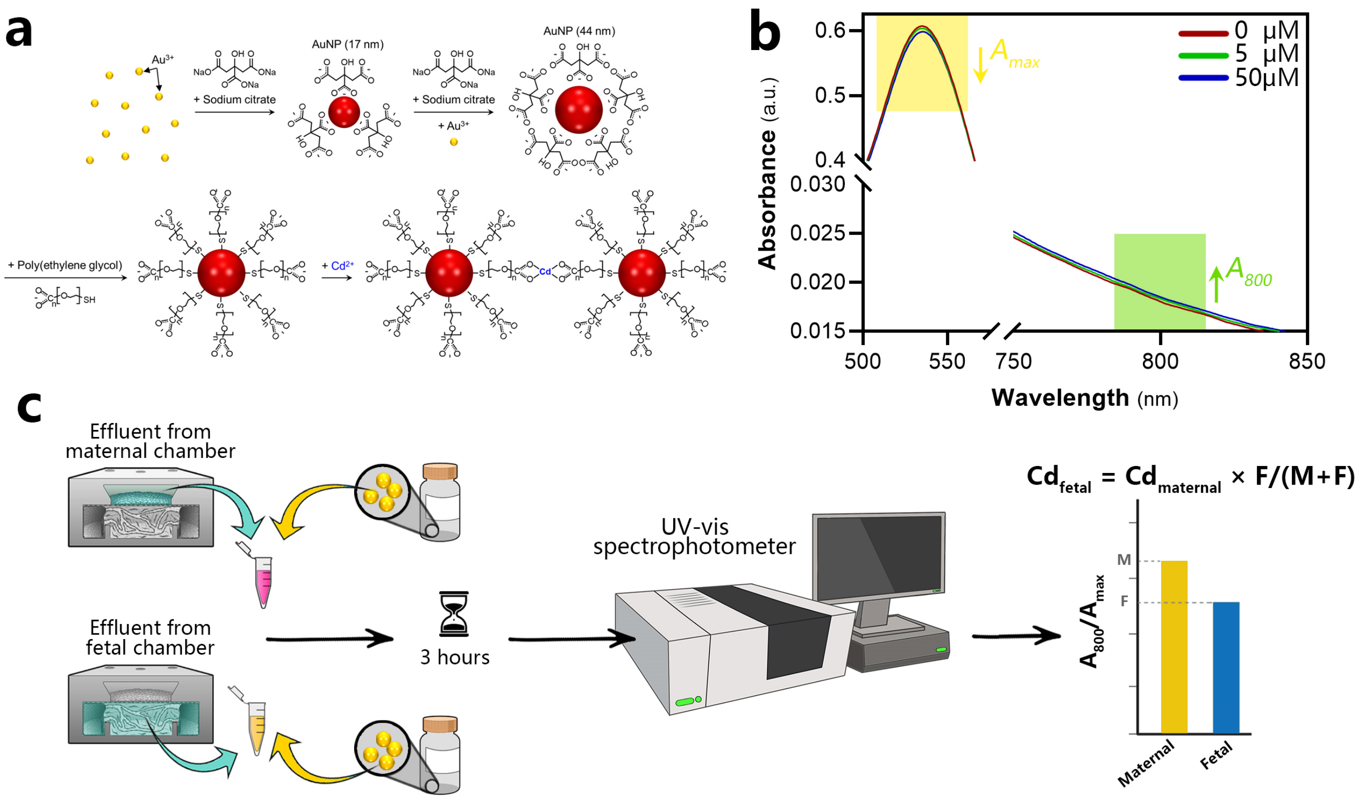

### Supplementary Figure 4. Heatmap of metabolites in the maternal compartment

A heatmap showing the relative abundance of the top 183 metabolites detected in all groups tested. Each column within a given group represents an independent replicate.

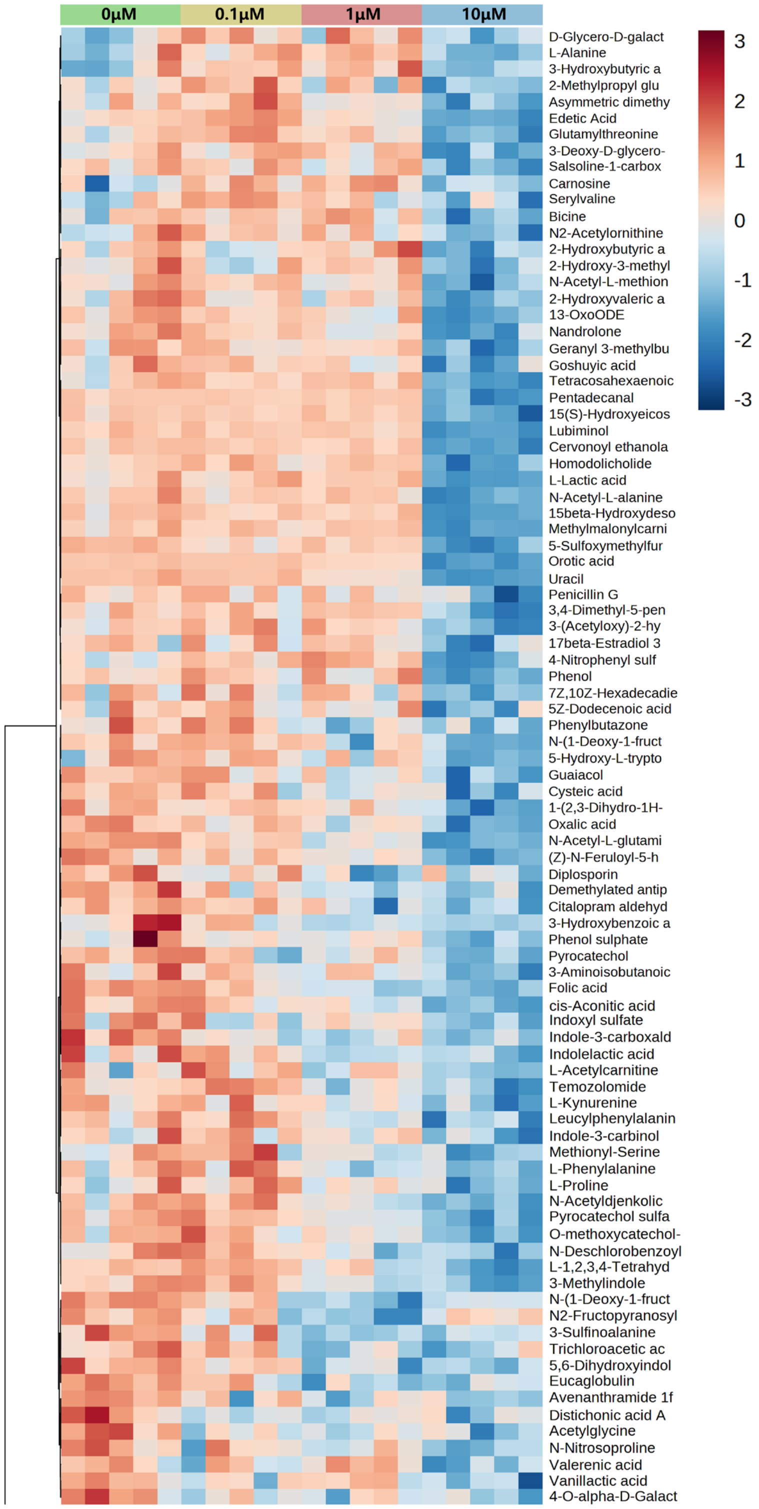

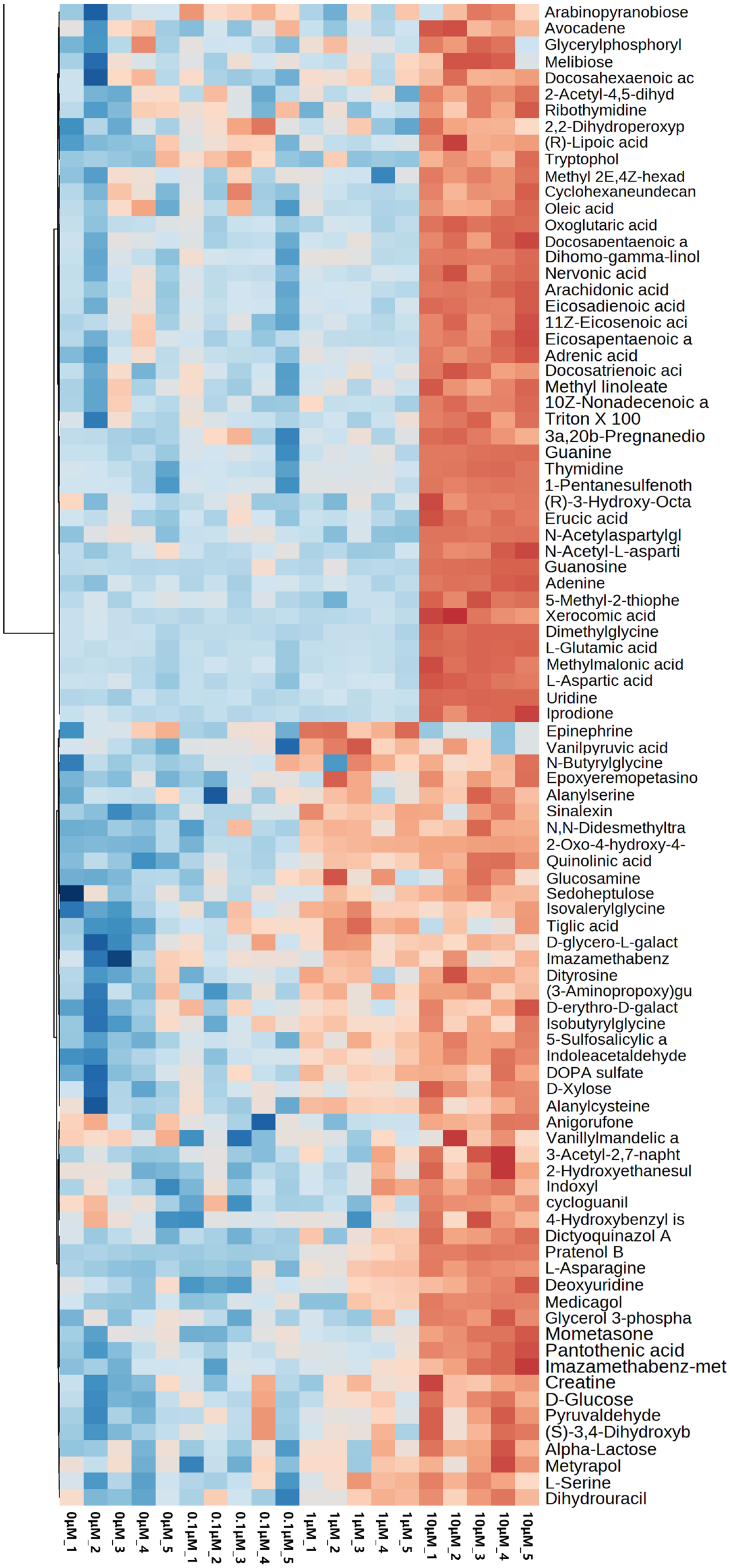
Arabinopyranobiose
Avocadene
Glycerylphosphoryl
Melibiose
Docosahexaenoic ac
2-Acetyl-4,5-dihyd
Ribothymidine
2,2-Dihydroperoxyp
(R)-Lipoic acid
Tryptophol
Methyl 2E,4Z-hexad
Cyclohexaneundecan
Oleic acid
Oxoglutaric acid
Docosapentaenoic a
Dihomo-gamma-linol
Nervonic acid
Arachidonic acid
Eicosadienoic acid
11Z-Eicosenoic aci
Eicosapentaenoic a
Adrenic acid
Docosatrienoic aci
Methyl linoleate
10Z-Nonadecenoic a
Triton X 100
3a,20b-Pregnanedio
Guanine
Thymidine
1-Pentanesulfenoth
(R)-3-Hydroxy-Octa
Erucic acid
N-Acetylaspartylgl
N-Acetyl-L-asparti
Guanosine
Adenine
5-Methyl-2-thiophe
Xerocomic acid
Dimethylglycine
L-Glutamic acid
Methylmalonic acid
L-Aspartic acid
Uridine
Iprodione
Epinephrine
Vanilpyruvic acid
N-Butyrylglycine
Epoxyeremopetasino
Alanylserine
Sinalexin
N,N-Didesmethyltra
2-Oxo-4-hydroxy-4-
Quinolinic acid
Glucosamine
Sedoheptulose
Isovalerylglycine
Tiglic acid
D-glycero-L-galact
Imazamethabenz
Dityrosine
(3-Aminopropoxy)gu
D-erythro-D-galact
Isobutyrylglycine
5-Sulfosalicylic a
Indoleacetaldehyde
DOPA sulfate
D-Xylose
Alanylcysteine
Anigorufone
Vanillylmandelic a
3-Acetyl-2,7-napht
2-Hydroxyethanesul
Indoxyl
cycloguanil
4-Hydroxybenzyl is
Dictyoquinazol A
Pratenol B
L-Asparagine
Deoxyuridine
Medicagol
Glycerol 3-phospha
Mometasone
Pantothenic acid
Imazamethabenz-met
Creatine
D-Glucose
Pyruvaldehyde
(S)-3,4-Dihydroxyb
Alpha-Lactose
Metyrapol
L-Serine
Dihydrouracil
0μM_1
0μM_2
0μM_3
0μM_4
0μM_5
0.1μM_1
0.1μM_2
0.1μM_3
0.1μM_4
0.1μM_5
1μM_1
1μM_2
1μM_3
1μM_4
1μM_5
10μM_1
10μM_2
10μM_3
10μM_4
10μM_5

### Supplementary Figure 5. Comparison of select metabolites in the maternal compartment

**a-b,** Volcano plots showing the top differentially regulated metabolites due to cadmium exposure at 10 μM (**a**) or 1 μM (**b**). **c-d,** Boxplots showing the normalized concentrations of select carbohydrate (**c**), nucleotide (**d**), peptide (**e**), and lipid (**f**) metabolites detected in the maternal compartment. Each box shows minimum, 25$^{th}$ percentile, mean, and 75$^{th}$ percentile. Ctrl represents an untreated negative control group. Data were normalized to the median. Data show mean ± SD with $n$ = 4 for each group. ns = not significant, *$P < 0.05$, **$P < 0.01$, ***$P < 0.001$, and **** $P < 0.0001$.

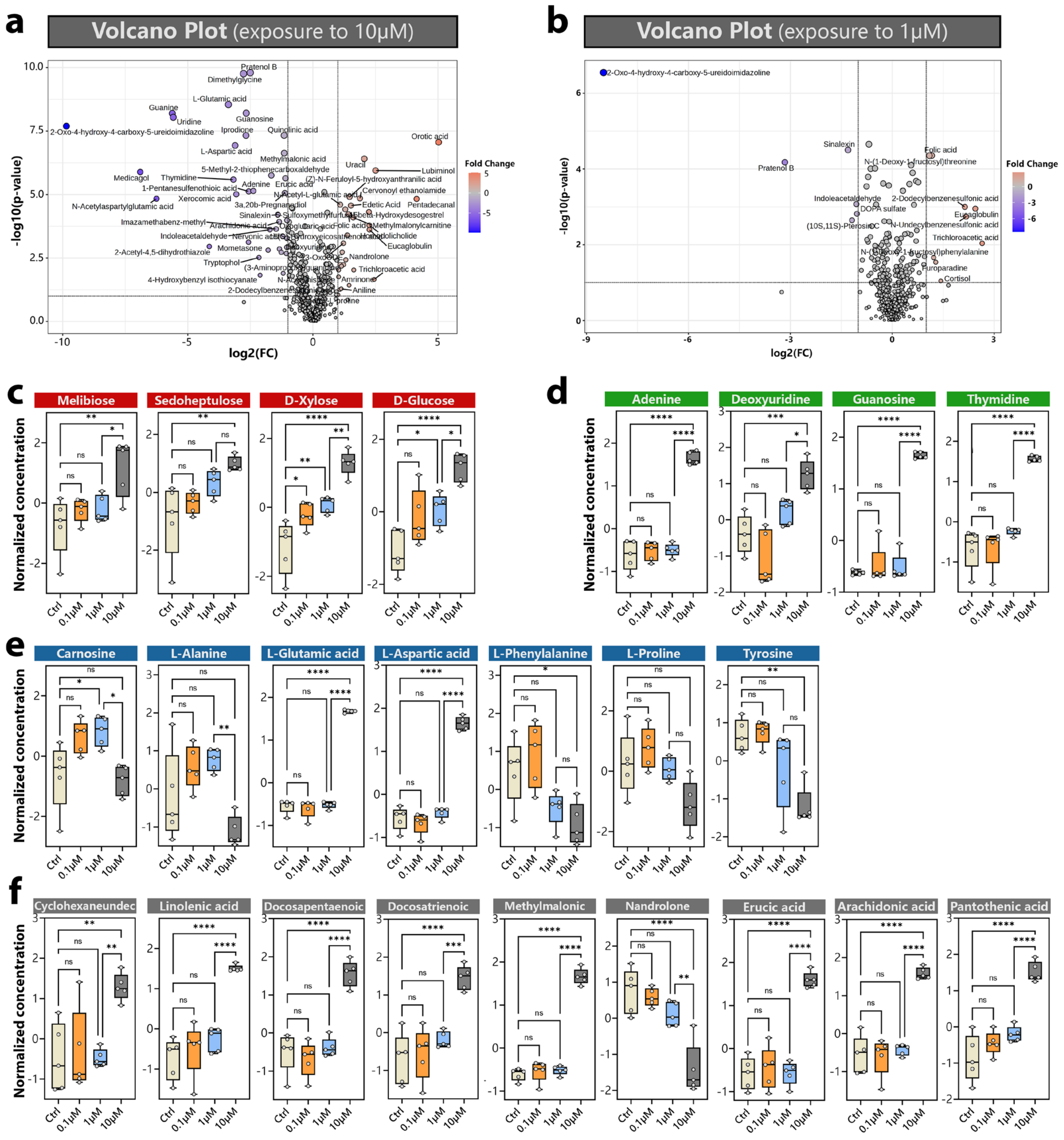

### Supplementary Figure 6. Heatmap of metabolites in the fetal compartment

A heatmap showing the relative abundance of top 255 metabolites detected in all groups tested. Each column within a given group represents an independent replicate.

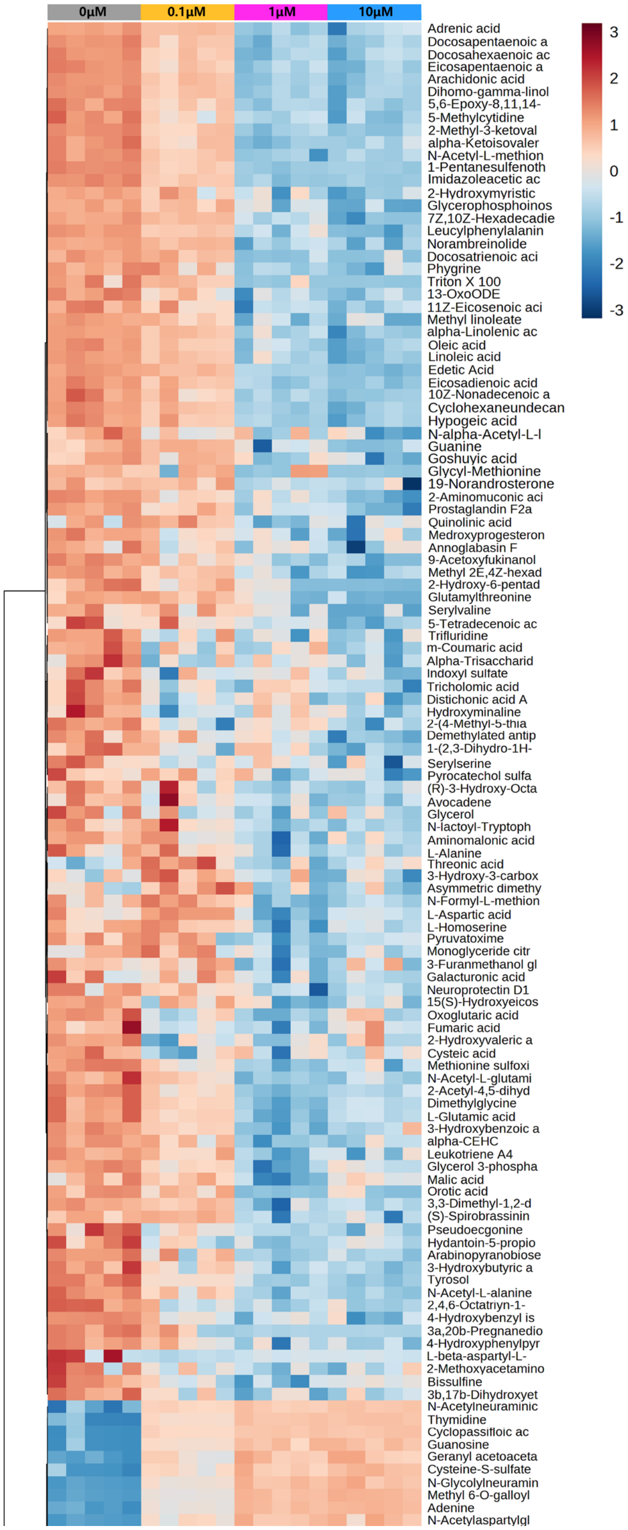

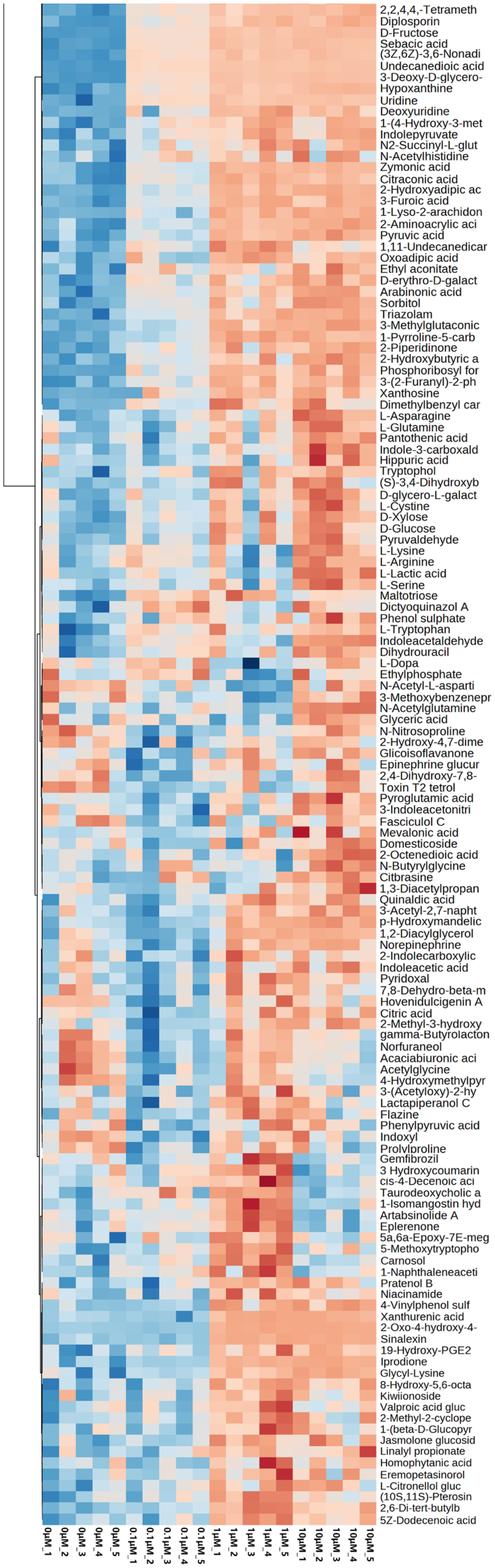

2,2,4,4,-Tetrameth
Diplosporin
D-Fructose
Sebacic acid
(3Z,6Z)-3,6-Nonadi
Undecanedioic acid
3-Deoxy-D-glycero-
Hypoxanthine
Uridine
Deoxyuridine
1-(4-Hydroxy-3-met
Indolepyruvate
N2-Succinyl-L-glut
N-Acetylhistidine
Zymonic acid
Citraconic acid
2-Hydroxyadipic ac
3-Furoic acid
1-Lyso-2-arachidon
2-Aminoacrylic aci
Pyruvic acid
1,11-Undecanedicar
Oxoadipic acid
Ethyl aconitate
D-erythro-D-galact
Arabinonic acid
Sorbitol
Triazolam
3-Methylglutaconic
1-Pyrroline-5-carb
2-Piperidinone
2-Hydroxybutyric a
Phosphoribosyl for
3-(2-Furanyl)-2-ph
Xanthosine
Dimethylbenzyl car
L-Asparagine
L-Glutamine
Pantothenic acid
Indole-3-carboxald
Hippuric acid
Tryptophol
(S)-3,4-Dihydroxyb
D-glycero-L-galact
L-Cystine
D-Xylose
D-Glucose
Pyruvaldehyde
L-Lysine
L-Arginine
L-Lactic acid
L-Serine
Maltotriose
Dictyoquinazol A
Phenol sulphate
L-Tryptophan
Indoleacetaldehyde
Dihydrouracil
L-Dopa
Ethylphosphate
N-Acetyl-L-asparti
3-Methoxybenzenepr
N-Acetylglutamine
Glyceric acid
N-Nitrosoproline
2-Hydroxy-4,7-dime
Glicoisoflavanone
Epinephrine glucur
2,4-Dihydroxy-7,8-
Toxin T2 tetrol
Pyroglutamic acid
3-Indoleacetonitri
Fasciculol C
Mevalonic acid
Domesticoside
2-Octenedioic acid
N-Butyrylglycine
Citbrasine
1,3-Diacetylpropan
Quinaldic acid
3-Acetyl-2,7-napht
p-Hydroxymandelic
1,2-Diacylglycerol
Norepinephrine
2-Indolecarboxylic
Indoleacetic acid
Pyridoxal
7,8-Dehydro-beta-m
Hovenidulcigenin A
Citric acid
2-Methyl-3-hydroxy
gamma-Butyrolacton
Norfuraneol
Acaciabiuronic aci
Acetylglycine
4-Hydroxymethylpyr
3-(Acetyloxy)-2-hy
Lactapiperanol C
Flazine
Phenylpyruvic acid
Indoxyl
Prolylproline
Gemfibrozil
3 Hydroxycoumarin
cis-4-Decenoic aci
Taurodeoxycholic a
1-Isomangostin hyd
Artabsinolide A
Eplerenone
5a,6a-Epoxy-7E-meg
5-Methoxytryptopho
Carnosol
1-Naphthaleneaceti
Pratenol B
Niacinamide
4-Vinylphenol sulf
Xanthurenic acid
2-Oxo-4-hydroxy-4-
Sinalexin
19-Hydroxy-PGE2
Iprodione
Glycyl-Lysine
8-Hydroxy-5,6-octa
Kiwiionoside
Valproic acid gluc
2-Methyl-2-cyclope
1-(beta-D-Glucopyr
Jasmolone glucosid
Linalyl propionate
Homophytanic acid
Eremopetasinorol
L-Citronellol gluc
(10S,11S)-Pterosin
2,6-Di-tert-butylb
5Z-Dodecenoic acid
0μM_1
0μM_2
0μM_3
0μM_4
0μM_5
0.1μM_1
0.1μM_2
0.1μM_3
0.1μM_4
0.1μM_5
1μM_1
1μM_2
1μM_3
1μM_4
1μM_5
10μM_1
10μM_2
10μM_3
10μM_4
10μM_5

### Supplementary Figure 7. Comparison of select metabolites in the fetal compartment.

**a-b,** Volcano plots showing top differentially regulated metabolites due to cadmium exposure at 10 μM (**a**) or 1 μM (**b**). **c-d,** Boxplots showing the normalized concentrations of select carbohydrate (**c**), nucleotide (**d**), peptide (**e**), and lipid (**f**) metabolites detected in the fetal compartment. Each box shows minimum, 25$^{th}$ percentile, mean, and 75$^{th}$ percentile. Ctrl represents an untreated negative control group. Data are normalized to the median. Data show mean ± SD with $n$ = 4 for each group. ns = not significant, *$P < 0.05$, **$P < 0.01$, ***$P < 0.001$, and **** $P < 0.0001$.

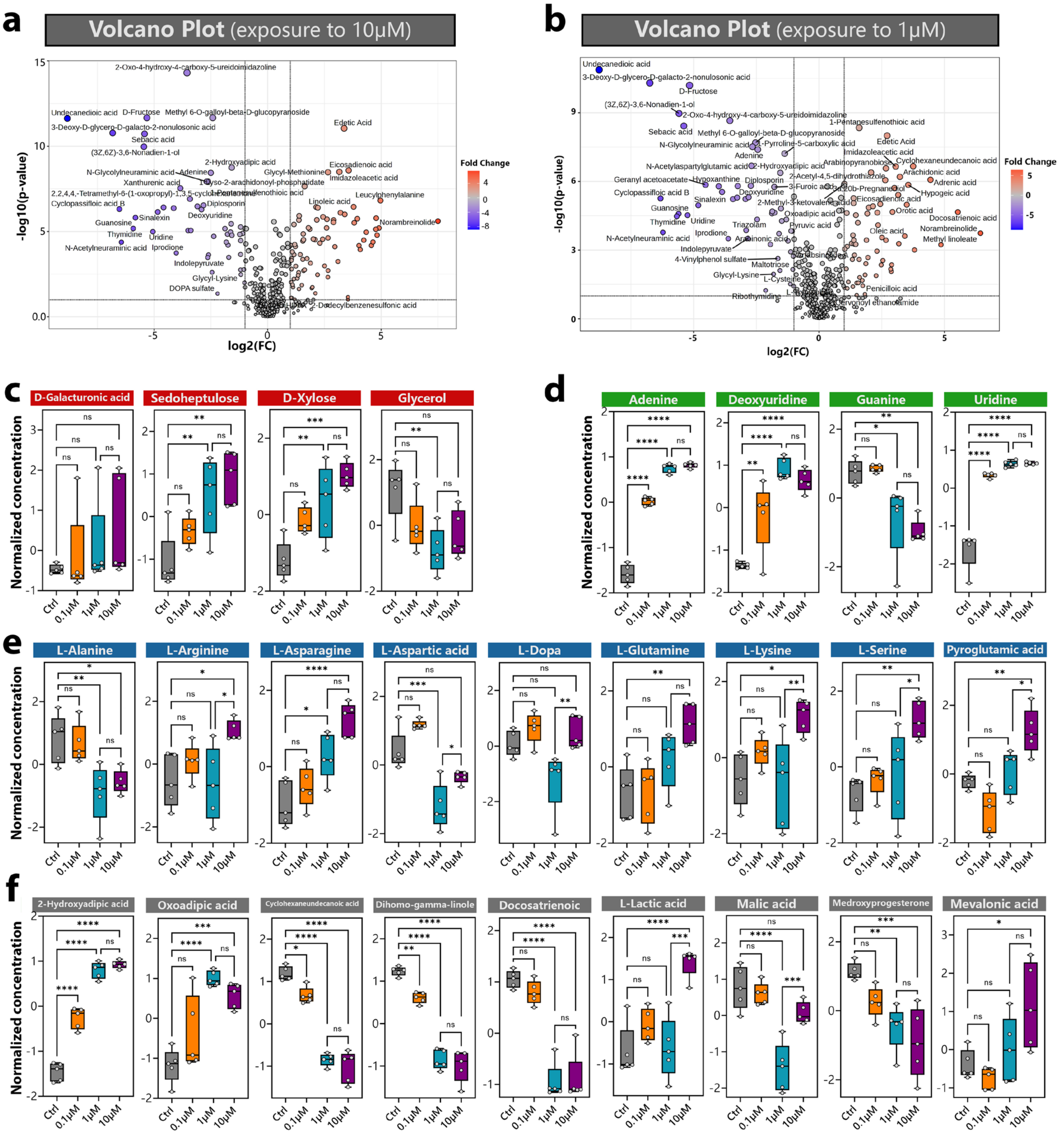

### Supplementary Figure 8. Comparison of cadmium-induced cytokine production in BeWo cell- and primary cell-based models

The BeWo cell-containing model produces higher levels of pro-inflammatory cytokines in both the maternal and fetal compartments than was measured in the primary cell-based model. The graphs were generated using data presented in **Figs. 3a, 3b,** and **4i**. Data show mean ± SD with $n$ = 4 for each group. ns = not significant, *$P < 0.05$*, **$P < 0.01$**, ***$P < 0.001$***, and **** $P < 0.0001$.

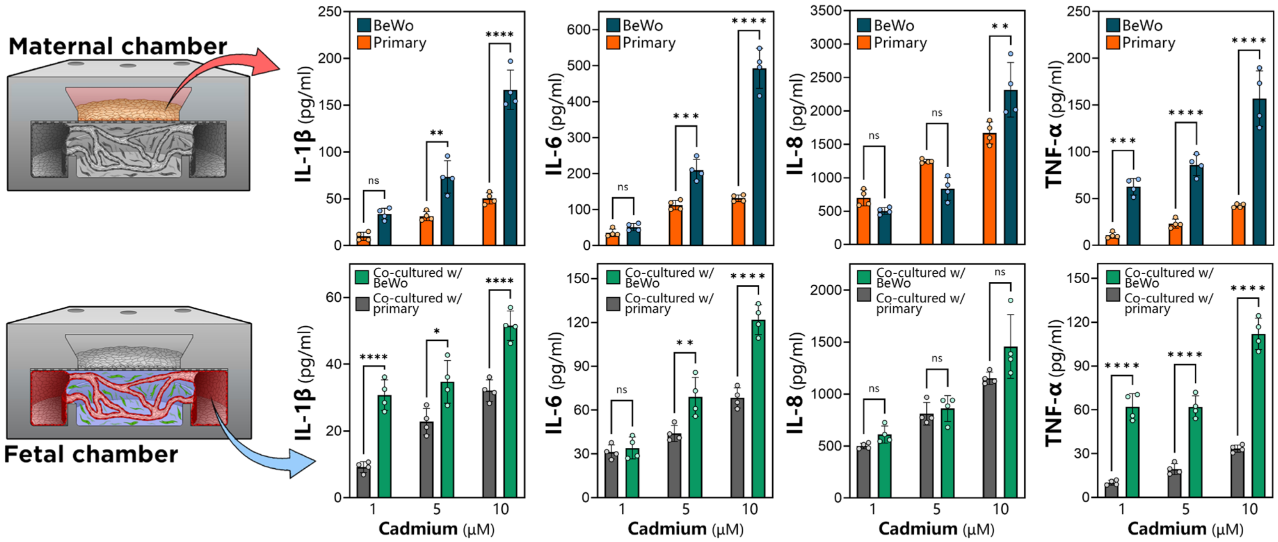

**Supplementary Table 1. Selected maternal metabolites and their association with human placental toxicity of environmental metals**

| No. | Metabolite | Type | KEGG Entry | Change | Association with placental toxicity of environmental metals | Toxicant | Ref. |
|---|---|---|---|---|---|---|---|
| 1 | Uridine | Nucleic acids | C00295 | Upregulated | Known | Arsenic | 1 |
| 2 | Uracil | Nucleic acids | C00106 | Downregulated | Known | Polychlorinated Biphenyls | 2 |
| 3 | L-Glutamic acid | Amino acids | C00025 | Upregulated | Known | Cadmium, mercury, cobalt, copper, thallium, and vanadium | 3,4 |
| 4 | Methylmalonic acid | Lipids | C02170 | Upregulated | **Unknown** | - | - |
| 5 | L-Aspartic acid | Amino acids | C00049 | Upregulated | Known | Tobacco (contains various metals) | 5,6 |
| 6 | Guanosine | Nucleic acids | C00387 | Upregulated | **Unknown** | - | - |
| 7 | Adenine | Nucleic acids | C00147 | Upregulated | Known | Cadmium | 7 |
| 8 | Pentadecanal | Lipids | C01948 | Downregulated | **Unknown** | - | - |
| 9 | Oxoglutaric acid | Organic acids | C00026 | Upregulated | **Unknown** | - | - |
| 10 | L-Asparagine | Amino acids | C00152 | Upregulated | Known | Tobacco (contains various metals) | 5,6 |
| 11 | Erucic acid | Lipids | C08316 | Upregulated | **Unknown** | - | - |
| 12 | Nervonic acid | Lipids | C08323 | Upregulated | **Unknown** | - | - |
| 13 | L-Lactic acid | Organic acids | C00186 | Downregulated | Known | Tobacco (contains various metals) | 6,8 |
| 14 | Thymidine | Nucleic acids | C00214 | Upregulated | **Unknown** | - | - |
| 15 | Arachidonic acid | Lipids | C00219 | Upregulated | Known | Particulate matter (PM2.5) (may contain metals) | 9,10 |
| 16 | Eicosadienoic acid | Lipids | C16525 | Upregulated | **Unknown** | - | - |
| 17 | Pantothenic acid | Vitamins and cofactors | C00864 | Upregulated | Known | Cadmium | 11 |
| 18 | Eicosapentaenoic acid | Lipids | C06428 | Upregulated | Known | Tobacco (contains various metals) and articulate matter (PM2.5) (may contain metals) | 5,12 |
| 19 | Docosapentaenoic acid (22n-6) | Lipids | C16513 | Upregulated | **Unknown** | - | - |
| 20 | Folic acid | Vitamins and cofactors | C00504 | Downregulated | **Unknown** | - | - |
| 21 | Adrenic acid | Lipids | C16527 | Upregulated | Known | Cadmium, chromium, manganese, nickel, copper, zinc, mercury, and lead | 13 |
| 22 | Dihomo-gamma-linolenic acid | Lipids | C03242 | Upregulated | **Unknown** | - | - |
| 23 | 13-OxoODE | Lipids | C14765 | Downregulated | Known | Particulate matter (PM2.5) (may contain metals) | 10 |
| 24 | D-Xylose | Carbohydrates | C00181 | Upregulated | **Unknown** | - | - |
| 25 | cis-Aconitic acid | Organic acids | C00417 | Downregulated | **Unknown** | - | - |
| 26 | Nandrolone | Lipids | C07254 | Downregulated | **Unknown** | - | - |
| 27 | Oxalic acid | Organic acids | C00209 | Downregulated | **Unknown** | - | - |
| 28 | Docosatrienoic acid | Lipids | C16534 | Upregulated | **Unknown** | - | - |
| 29 | Deoxyuridine | Nucleic acids | C00526 | Upregulated | **Unknown** | - | - |
| 30 | Tiglic acid | Lipids | C08279 | Upregulated | **Unknown** | - | - |
| 31 | D-Glucose | Carbohydrates | C00031 | Upregulated | **Unknown** | - | - |
| 32 | L-Serine | Peptides | C00065 | Upregulated | Known | Tobacco (contains various metals) | 6,14 |
| 33 | (R)-Lipoic acid | Lipids | C16241 | Upregulated | **Unknown** | - | - |
| 34 | N-(1-Deoxy-1-fructosyl)tyrosine | Peptides | C00082 | Downregulated | Known | Cadmium and tobacco (contains various metals) | 6,15 |

| | | | | | | | |
|---|---|---|---|---|---|---|---|
| 35 | 2-Hydroxybutyric acid | Lipids | C05984 | Downregulated | **Unknown** | - | - |
| 36 | 3-Sulfinoalanine | Peptides | C00606 | Downregulated | **Unknown** | - | - |
| 37 | Glucosamine | Carbohydrates | C00329 | Upregulated | Known | Polychlorinated Bi-phenyls and particu-late matter (PM2.5) (may contain metals) | 2,10 |
| 38 | Cyclohexaneun-decanoic acid | Lipids | C12100 | Upregulated | **Unknown** | - | - |
| 39 | L-Alanine | Peptides | C00041 | Downregulated | **Unknown** | - | - |
| 40 | L-Phenylalanine | Peptides | C00079 | Downregulated | **Unknown** | - | - |
| 41 | Epinephrine | Hormones and transmitters | C00547 | Upregulated | **Unknown** | - | - |
| 42 | Carnosine | Peptides | C00386 | Upregulated | **Unknown** | - | - |
| 43 | Oleic acid | Lipids | C00712 | Upregulated | **Unknown** | - | - |
| 44 | Sedoheptulose | Carbohydrates | C02076 | Upregulated | Known | Polychlorinated Bi-phenyls | 2 |
| 45 | Melibiose | Carbohydrates | C05402 | Upregulated | **Unknown** | - | - |
| 46 | Penicillin G | Antibiotics | C05551 | Downregulated | **Unknown** | - | - |
| 47 | Trichloroacetic acid | Peptides | C11150 | Downregulated | **Unknown** | - | - |
| 48 | 3-Hydroxybutyric acid | Organic acids | C01089 | Downregulated | Known | Cadmium | 16,17 |
| 49 | L-Proline | Peptides | C00148 | Downregulated | Known | Cadmium | 18,19 |
| 50 | Docosahexaenoic acid | Lipids | C06429 | Upregulated | **Unknown** | - | - |
| 51 | Valerenic acid | Lipids | C09743 | Downregulated | **Unknown** | - | - |
| 52 | 17beta-Estradiol 3-sulfate | Lipids | C09743 | Downregulated | **Unknown** | - | - |

### Supplementary Table 2. Selected fetal metabolites and their association with human placental toxicity of environmental metals

| No. | Metabolite | Type | KEGG Entry | Change | Association with placental toxicity of environmental metals | Toxicant | Ref. |
|---|---|---|---|---|---|---|---|
| 1 | D-Fructose | Carbohydrates | C00095 | Upregulated | **Unknown** | - | - |
| 2 | Sebacic acid | Lipids | C08277 | Upregulated | **Unknown** | - | - |
| 3 | Edetic Acid | Vitamins and co-factors | C00284 | Downregulated | **Unknown** | - | - |
| 4 | Adenine | Nucleic acids | C00147 | Upregulated | Known | Cadmium | 7,20 |
| 5 | 2-Hydroxyadipic acid | Lipids | C02360 | Upregulated | **Unknown** | - | - |
| 6 | Eicosadienoic acid | Lipids | C16525 | Downregulated | **Unknown** | - | 21 |
| 7 | Arachidonic acid | Lipids | C00219 | Downregulated | Known | Particulate matter (PM2.5) (may contain metals) | 9,10 |
| 8 | Cyclohexaneun-decanoic acid | Lipids | C12100 | Downregulated | **Unknown** | - | - |
| 9 | Dihomo-gamma-lin-olenic acid | Lipids | C03242 | Downregulated | **Unknown** | - | - |
| 10 | Guanosine | Nucleic acids | C00387 | Upregulated | **Unknown** | - | - |
| 11 | Uridine | Nucleic acids | C00299 | Upregulated | Known | Arsenic | 1,22 |
| 12 | Thymidine | Nucleic acids | C00214 | Upregulated | **Unknown** | - | - |
| 13 | Citraconic acid | Lipids | C02226 | Upregulated | Known | Polychlorinated Bi-phenyls | 2 |
| 14 | L-Glutamic acid | Peptides | C00025 | Downregulated | Known | Cadmium, mercury, cobalt, copper, thal-lium, and vanadium and tobacco (contains various metals) | 3,4,2 3 |
| 15 | Oleic acid | Lipids | C00712 | Downregulated | **Unknown** | - | - |
| 16 | Linoleic acid | Lipids | C01595 | Downregulated | Known | Cadmium | 11,14 |
| 17 | Docosahexaenoic acid | Lipids | C06429 | Downregulated | **Unknown** | - | - |
| 18 | N-Acetylneuraminic acid | Carbohydrates | C00270 | Upregulated | **Unknown** | - | - |
| 19 | Pyruvic acid | Lipids | C00022 | Upregulated | Known | Inorganic arsenic (iAs) | 24 |
| 20 | Docosatrienoic acid | Lipids | C16534 | Downregulated | **Unknown** | - | - |
| 21 | alpha-Linolenic acid | Lipids | C06427 | Downregulated | **Unknown** | - | - |
| 22 | 2-Hydroxybutyric acid | Lipids | C05984 | Upregulated | **Unknown** | - | - |
| 23 | Adrenic acid | Lipids | C16527 | Downregulated | Known | Cadmium, chromium, manganese, nickel, copper, zinc, mercury, and lead | 13 |
| 24 | D-Sorbitol | Carbohydrates | C00794 | Upregulated | **Unknown** | - | - |
| 25 | Prostaglandin F2a | Lipids | C00639 | Downregulated | **Unknown** | - | - |
| 26 | Glycerophospho-inositol | Lipids | C03819 | Downregulated | Known | Tobacco (contains various metals) | 6 |
| 27 | L-Aspartic acid | Peptides | C00049 | Downregulated | Known | Tobacco (contains various metals) | 5,6 |
| 28 | Deoxyuridine | Nucleic acids | C00526 | Upregulated | **Unknown** | - | - |
| 29 | 13-OxoODE | Lipids | C14765 | Downregulated | Known | Particulate matter (PM2.5) (may contain metals) | 10 |
| 30 | Malic acid | Organic acids | C00149 | Downregulated | **Unknown** | - | - |
| 31 | 3-Hydroxybutyric acid | Lipids | C01089 | Downregulated | Known | Cadmium | 16,17 |
| 32 | L-Lactic acid | Organic acids | C00186 | Upregulated | Known | Tobacco (contains various metals) | 6,8 |
| 33 | N-Formyl-L-methio-nine | Peptides | C03145 | Downregulated | Known | Arsenic, tobacco (contains various | 1,23, 24 |

| | | | | | | | |
|---|---|---|---|---|---|---|---|
| | | | | | | metals) and inorganic arsenic (iAs) | |
| 34 | Oxoadipic acid | Lipids | C00322 | Upregulated | **Unknown** | - | - |
| 35 | L-Asparagine | Peptides | C00152 | Upregulated | Known | Tobacco (contains various metals) | 5,6 |
| 36 | Leukotriene | Lipids | C00909 | | Known | Tobacco (contains various metals) | 5 |
| 37 | Oxoglutaric acid | Organic acids | C00026 | Downregulated | **Unknown** | - | - |
| 38 | Norepinephrine | Hormones and transmitters | C00547 | Upregulated | Known | Polychlorinated Bi-phenyls | 2 |
| 39 | Pyroglutamic acid | Peptides | C01879 | Upregulated | **Unknown** | - | - |
| 40 | L-Homoserine | Peptides | C00263 | Downregulated | Known | Tobacco (contains various metals) | 25 |
| 41 | D-Xylose | Carbohydrates | C00181 | Upregulated | **Unknown** | - | - |
| 42 | Guanine | Nucleic acids | C00242 | Downregulated | Known | Cadmium | 26 |
| 43 | Citric acid | Organic acids | C00158 | Upregulated | Known | Phthalates and metals | 27 |
| 44 | Medroxyprogester-one | Lipids | C07119 | Downregulated | **Unknown** | - | - |
| 45 | D-Glucose | Carbohydrates | C00031 | Upregulated | **Unknown** | - | - |
| 46 | Pyridoxal | Vitamins & co-factors | C00250 | Upregulated | **Unknown** | - | - |
| 47 | L-Alanine | Peptides | C00041 | Downregulated | **Unknown** | - | - |
| 48 | L-Glutamine | Peptides | C00064 | Upregulated | Known | Tobacco (contains various metals) | 7,23 |
| 50 | L-Serine | Peptides | C00065 | Upregulated | Known | Tobacco (contains various metals) | 6,14 |
| 51 | Taurodeoxycholic acid | Lipids | C05463 | Downregulated | Known | Persistent organic pollutants (POPs) | 28 |
| 52 | L-Dopa | Peptides | C00355 | Downregulated | **Unknown** | - | - |
| 53 | L-Tryptophan | Peptides | C00078 | Upregulated | Known | Tobacco (contains various metals) | 23 |
| 54 | gamma-Butyrolac-tone | Lipids | C01770 | Downregulated | **Unknown** | - | - |
| 55 | L-Lysine | Peptides | C00047 | Upregulated | Known | Cadmium, cobalt, copper, cesium, man-ganese, thallium and vanadium | 4,19 |
| 56 | Mevalonic acid | Lipids | C00418 | Upregulated | **Unknown** | - | - |
| 57 | Niacinamide | Vitamins and co-factors | C00153 | Upregulated | Known | Particulate matter (PM2.5) (may contain metals) | 10 |
| 58 | Pantothenic acid | Vitamins and co-factors | C00864 | Upregulated | Known | Cadmium and zinc | 11 |
| 59 | Glycerol | Carbohydrates | C00116 | Downregulated | Known | Inorganic arsenic (iAs) | 24 |
| 60 | D-Galacturonic acid | Carbohydrates | C00333 | Upregulated | Known | Cadmium and particu-late matter (PM2.5) (may contain metals) | 10,29 |
| 61 | L-Arginine | Peptides | C00062 | Upregulated | Known | Tobacco (contains various metals) | 23,30 |

### Supplementary Table 3. Key Resources

| Reagent or resource | Source | Identifier |
|---|---|---|
| **Antibodies** | | |
| Alexa Fluor 488 Anti-CD31 | abcam | ab215911 |
| Anti-E-Cadherin | abcam | ab40772 |
| Anti-E-Cadherin | abcam | ab1416 |
| Anti-Glucose Transporter (GLUT1) | abcam | ab115730 |
| Anti-CD163 | abcam | ab182422 |
| Anti-CD163 | Invitrogen | MA5-17716 |
| Anti-ZO-1 | Invitrogen | 61-7300 |
| Anti-ZO-1 | Invitrogen | 33-9100 |
| Alexa Fluor 488 Anti-Fibronectin | Invitrogen | 53-9869-82 |
| Anti-alpha -Smooth Muscle | Invitrogen | 14-9760-82 |
| Anti-alpha -Smooth Muscle | R&D Systems | MAB1420 |
| Anti-BCRP | Santa Cruz | Sc-18841 |
| Alexa Fluor™ 488 Phalloidin | Thermofisher | A-12379 |
| Alexa Fluor™ 647 Phalloidin | Thermofisher | A-22287 |
| Donkey anti-mouse Secondary Antibody Alexa Fluor™ Plus 555 | Invitrogen | A-32773 |
| Goat anti-mouse Secondary Antibody Alexa Fluor™ Plus 488 | Invitrogen | A-32723 |
| Goat anti-mouse Secondary Antibody Alexa Fluor™ Plus 647 | Invitrogen | A-32728 |
| Goat anti-rabbit Secondary Antibody Alexa Fluor™ Plus 555 | Invitrogen | A-32732 |
| Donkey anti-rabbit Secondary Antibody Alexa Fluor™ 488 | Invitrogen | A-32790 |
| **Chemicals, peptides, and recombinant proteins** | | |
| DMEM/F-12, HEPES, no phenol red | Gibco/Invitrogen | Catalog # 11039021 |
| EGM™-2 MV BulletKit™ | Lonza | Catalog # CC-3202 |
| EGM™-2 BulletKit™ | Lonza | Catalog # CC-3162 |
| FGM™-2 BulletKit™ | Lonza | Catalog # CC-3132 |
| CTBPRO2GRO™ trophoblast media | Amnion Foundation | Catalog # CTBPRO2GRO |
| Invitrogen™ Lucifer Yellow CH, Lithium Salt, 25 mg | Invitrogen | Catalog # L453 |
| Cadmium chloride solution | Sigma Aldrich | Catalog # 21115 |
| EasySep™ Magnet | STEMCELL Technologies | Catalog # 18000 |
| EasySep™ Direct Human Neutrophil Isolation Kit | STEMCELL Technologies | Catalog # 100-0404 |
| **Assays** | | |
| Human IL-6 ELISA kit | Sigma Aldrich | RAB0306 |
| Human IL-8 / CXCL8 ELISA Kit | Sigma Aldrich | RAB0319 |
| Human TNF alpha ELISA Kit | abcam | Ab181421 |
| Human IL-1β ELISA Kit | Sigma Aldrich | RAB0273 |
| Human TGF-beta 1 DueSet ELISA | R&D Systems | DY240 |
| β-HCG ELISA | DiaMetra | DKO014 |
| Cytotoxicity Detection Kit$^{PLUS}$ (LDH) | Roche | 04744926001 |
| **Cells** | | |
| Human Cytotrophoblast Cells | Amnion Foundation | Cat #1230 |
| Placental Stromal Cells/Fibroblasts | Amnion Foundation | Cat# 1250 |
| Germinal-Origin Microvascular Endothelial Cells | Amnion Foundation | Cat #1245 |
| Hofbauer Cells | Amnion Foundation | Cat #1220 |
| BeWo-b30 human choriocarcinoma cells | Donated by Dr. Lauren M. Aleksunes (Rutgers University) | |

| | | |
|---|---|---|
| **Oligonucleotides** | | |
| Primers for RT PCR | This paper | see **Supplementary Table 4** |

### Supplementary Table 4. List of primers

| Gene | Forward | Reverse |
|---|---|---|
| **GAPDH** | *CGCTCTCTGCTCCTCCTGTT* | *CCATGGTGTCTGAGCGATGT* |
| **Syn-1** | *GCAACCACGAACGGACATC* | *GTATCCAAGACTCCACTCCAGC* |
| **Syn-2** | *CGGATACCTTCCCTAGTGCC* | *AGCTGAGGTTGCTGGTTCTG* |
| **hCG-α** | *CAGAATGCACGCTACAGGAA* | *CGTGTGGTTCTCCACTTTGA* |
| **hCG-β** | *GCACCAAGGATGGAGATGTT* | *GCACATTGACAGCTGAGAGC* |
| **ABCG2** | *GGATGAGCCTACAACTGGCTT* | *CTTCCTGAGGCCAATAAGGTG* |
| **ABCB1** | *GTGGTGGGAACTTTGGCTG* | *TACCTGGTCATGTCTTCCTCC* |
| **ABCB4** | *ATCGAGACGTTACCCCACAA* | *CATTCTGGATGGTGGACAGG* |
| **ABCC1** | *GTGTTTCTGGTCAGCCCAACT* | *TTGGATCTCAGGATGGCTAGG* |
| **ABCC2** | *TCCAACTGTGCTTCAAGC* | *GGCATCCACAGACATCAG* |
| **SLC2A1** | *GATGATGCGGGAGAAGAAGGT* | *ACAGCGTTGATGCCAGACAG* |
| **SLC2A3** | *CTTCCCCTCCGCTGCTCACTA* | *CAAAAGTCCTGCCACGGGTCT* |
| **SLC11A1** | *TGCATCTTGCTGAAGTATGTCACC* | *CTCCACCATCAGCCACAGGAT* |
| **Twist1** | *TCTCGGTCTGGAGGATGGA* | *CAATGACATCTAGGTCTCCG* |
| **PIGF** | *GGCTGTTCCCTTGCTTCCT* | *TACCACTTCCACCTCTGACGA* |
| **SDC1** | *GGATGACTCTGACAACTTCTCC* | *CTACAGCCTCTCCCTCCTT* |
| **ZIP8** | *CAGTGTGGTATCTCTACAGGATGGA* | *CAGTTTGGGCCCCTTCAAA* |
| **ZIP14** | *CAAGTCTGCAGTGGTGTTTG* | *GTGTCCATGATGATGCTCATTT* |
| **E-cad** | *GCCGAGAGCTACACGTTCAC* | *GTCGAGGGAAAAATAGGCTG* |
| **ZO-1** | *CAACATACAGTGACGCTTCACA* | *CACTATTGACGTTTCCCCACTC* |
| **Snail** | *AAGATGCACATCCGAAGCCA* | *CATTCGGGAGAAGGTCCGAG* |
| **Zeb1** | *TGCACTGAGTGTGGAAAAGC* | *TGGTGATGCTGAAAGAGACG* |
| **Zeb2** | *CGCTTGACATCACTGAAGGA* | *CTTGCCACACTCTGTGCATT* |
| **P-53** | *CATGAGCGCTGCTCAGATAG* | *ACACGCAAATTTCCTTCCAC* |
| **MT1A** | *CTTGGGATCTCCAACCTCAC* | *AGGAGCAGCAGCTCTTCTTG* |
| **MT2A** | *CCGACTCTAGCCGCCTCTT* | *GTGGAAGTCGCGTTCTTTACA* |
| **ZnT1** | *CCTGGGCTTCTTCTCTAGATTG* | *TTGTCTTGGAAAGGTTGTTCTG* |
| **ZnT2** | *CCTGGTCTCTGTACTGTCCATCT* | *GATCACGAACAGCTGTGAAGTC* |
| **SOD** | *GCAGAAGGCAAGCGGTGAAC* | *TAGCAGGACAGCAGATGAGT* |
| **CAT** | *GCGAATGGAGAGGCAGTGTAC* | *GAGTGACGTTGTCTTCATTAGCACTG* |
| **GPX1** | *AGATGTCATTCCTGCACACG* | *AAGGAGAAGCTTCCTCAGCC* |
| **BCL-2** | *TCCCTCGCTGCACAAATACTC* | *TTCTGCCCCTGCCAAATCT* |
| **Nrf2** | *CGCAGACATTCCCGTTTGTAGA* | *GTGACCGGGAATATCAGGAACAAG* |
| **NF-κB** | *GCAAAGGGAACATTCCGATAT* | *GCGACATCACATGGAAATCTA* |
| **TRPV2** | *TGTAGCCCTGGTGAGCCT* | *CCAACGGTCAGCATCACA* |
| **TRPV4** | *CTACGGCACCTATCGTCACC* | *CTGCGGCTGCTTCTCTATGA* |
| **Caspase3** | *AATTGTGGAATTGATGCGTGATG* | *CTACAACGATCCCCTCTGAAAAA* |
| **Caspase7** | *CCAATAAAGGATTTGACAGCC* | *GCATCTGTGTCATTGATGGG* |
| **Caspase9** | *ATGGACGAAGCGGATCGG* | *CCCTGGCCTTATGATGTT* |
| **VEGFR1** | *CAGGCCCAGTTTCTGCCATT* | *TTCCAGCTCAGCGTGGTCGTA* |
| **VEGFR2** | *CCAGCAAAAGCAGGGAGTCTGT* | *TGTCTGTGTCATCGGAGTGATATCC* |
| **PECAM1** | *GAGTATTACTGCACAGCCTTCA* | *AACCACTGCAATAAGTCCTTTC* |
| **CLDN5** | *GTTCGCCAACATTGTCGTCC* | *GTAGTTCTTCTTGTCGTAGTCGC* |
| **VCAM-1** | *CCGTCTCATTGACTTGCAGC* | *GATGTGGTCCCCTCATTCGT* |
| **COL1A1** | *ATCAACCGGAGGAATTTCCGT* | *CACCAGGACGACCAGGTTTTC* |
| **COL1A2** | *GGCCCTCAAGGTTTCCAAGG* | *CACCCTGTGGTCCAACAACTC* |
| **FN1** | *CAGGATCACTTACGGAGAAACAG* | *GCCAGTGACAGCATACACAGTG* |
| **CTGF** | *GCAGGCTAGAGAAGCAGAGC* | *ATGTCTTCATGCTGGTGCAG* |
| **PDGF** | *TGATCTCCAACGCCTGCT* | *TCATGTTCAGGTCCAACTCG* |
| **ICAM-1** | *AACCAGAGCCAGGAGACACT* | *GAGACCTCTGGCTTCGTCAG* |
| **MMP-2** | *ACATCAAGGGCATTCAGGAG* | *GCCTCCGTATACCGCATCAAT* |
| **MMP-9** | *GGGAAGATGCTGCTGTTCA* | *TCAACTCACTCCGGGAACTC* |
| **TIMP-1** | *TCAACCAGACCACCTTATACCA* | *ATCCGCAGACACTCCAT* |
